# Acoustic regularities in infant-directed speech and song across cultures

**DOI:** 10.1101/2020.04.09.032995

**Authors:** Courtney B. Hilton, Cody J. Moser, Mila Bertolo, Harry Lee-Rubin, Dorsa Amir, Constance M. Bainbridge, Jan Simson, Dean Knox, Luke Glowacki, Elias Alemu, Andrzej Galbarczyk, Grazyna Jasienska, Cody T. Ross, Mary Beth Neff, Alia Martin, Laura K. Cirelli, Sandra E. Trehub, Jinqi Song, Minju Kim, Adena Schachner, Tom A. Vardy, Quentin D. Atkinson, Amanda Salenius, Jannik Andelin, Jan Antfolk, Purnima Madhivanan, Anand Siddaiah, Caitlyn D. Placek, Gul Deniz Salali, Sarai Keestra, Manvir Singh, Scott A. Collins, John Q. Patton, Camila Scaff, Jonathan Stieglitz, Silvia Ccari Cutipa, Cristina Moya, Rohan R. Sagar, Mariamu Anyawire, Audax Mabulla, Brian M. Wood, Max M. Krasnow, Samuel A. Mehr

**Affiliations:** Department of Psychology, Harvard University, Cambridge, MA 02138, USA; Department of Cognitive and Information Sciences, University of California Merced, Merced, CA 95343, USA; Boston College Department of Psychology, Chestnut Hill, MA 02467, USA; Department of Communication, University of California Los Angeles, Los Angeles, CA 90095, USA; Department of Psychology, University of Amsterdam, 1012 WX Amsterdam, The Netherlands; Operations, Information, and Decisions Department, the Wharton School of the University of Pennsylvania, Philadelphia, PA 19104, USA; Department of Anthropology, Boston University, Boston, MA 02215, USA; Jinka University, Jinka, South Omo Zone, Ethiopia; Department of Environmental Health, Faculty of Health Sciences, Jagiellonian University Medical College, 31-066 Krakow, Poland; Department of Human Behavior, Ecology and Culture, Max Planck Institute for Evolutionary Anthropology, 04103 Leipzig, Germany; School of Psychology, Victoria University of Wellington, Wellington 6012, New Zealand; Department of Philosophy, Classics, History of Art and Ideas, University of Oslo, Oslo 0315, Norway; Department of Psychology, University of Toronto Scarborough, Toronto, Ontario M1C 1A4, Canada; Department of Psychology, University of Toronto Mississauga, Mississauga, Ontario L5L 1C6, Canada; Department of Mathematics, University of California Los Angeles, Los Angeles, CA 90095, USA; Department of Psychology, University of California, San Diego, La Jolla, CA 92093-0109, USA; School of Psychology, University of Auckland, Auckland 1010, New Zealand; Department of Linguistic and Cultural Evolution, Max Planck Institute for Evolutionary Anthropology, 04103 Leipzig, Germany; Department of Psychology, Åbo Akademi, 20500 Turku, Finland; Department of Health Promotion Sciences, College of Public Health, University of Arizona, Tucson, AZ 85724, USA; Department of Medicine, Division of Infectious Diseases, College of Medicine, University of Arizona, Tucson, AZ 85724, USA; Department of Family & Community Medicine, College of Medicine, University of Arizona, Tucson, AZ 85724, USA; Public Health Research Institute of India, Mysuru 570020, India; Department of Anthropology, Ball State University, Muncie, IN 47306, USA; Department of Anthropology, University College London, WC1H 0BW London, UK; Department of Human Evolutionary Biology, Harvard University, Cambridge, MA 02138, USA; Institute for Advanced Study in Toulouse, 31080 Toulouse Cedex 6, France; School of Human Evolution and Social Change, Arizona State University, Tempe, AZ 85281, USA; Division of Anthropology, California State University, Fullerton, CA 92831, USA; Institute of Evolutionary Medicine, University of Zurich, 8006 Zürich, Switzerland; Université Toulouse 1 Capitole, 31080 Toulouse Cedex 6, France; Universidad Nacional del Altiplano Puno, Puno 21001, Peru; Department of Anthropology, University of California, Davis, Davis, CA 95616, USA; Centre for Culture & Evolution, Brunel University London, UB8 3PH Uxbridge, UK; Future Generations University, Circle Ville, WV 26807, USA; Harpy Eagle Music Foundation, Georgetown, Guyana; Mang’ola, Tanzania; Department of Archaeology and Heritage, University of Dar es Salaam, Dar es Salaam, Tanzania; Department of Anthropology, University of California, Los Angeles, Los Angeles, CA 90095, USA; Division of Continuing Education, Harvard University, Cambridge, MA 02138, USA; Data Science Initiative, Harvard University, Cambridge, MA 02138, USA

## Abstract

The forms of many species’ vocal signals are shaped by their functions^1–15^. In humans, a salient context of vocal signaling is infant care, as human infants are altricial^16, 17^. Humans often alter their vocalizations to produce “parentese”, speech and song produced for infants that differ acoustically from ordinary speech and song^18–35^ in fashions that have been proposed to support parent-infant communication and infant language learning^36–39^; modulate infant affect^33, 40–45^; and/or coordinate communicative interactions with infants^46–48^. These theories predict a form-function link in infant-directed vocalizations, with consistent acoustic differences between infant-directed and adult-directed vocalizations across cultures. Some evidence supports this prediction^23, 27, 28, 32, 49–52^, but the limited generalizability of individual ethnographic reports and laboratory experiments^53^ and small stimulus sets^54^, along with intriguing reports of counterexamples^55–62^, leave the question open. Here, we show that people alter the acoustic forms of their vocalizations in a consistent fashion across cultures when speaking or singing to infants. We collected 1,615 recordings of infant- and adult-directed singing and speech produced by 410 people living in 21 urban, rural, and small-scale societies, and analyzed their acoustic forms. We found cross-culturally robust regularities in the acoustics of infant-directed vocalizations, such that infant-directed speech and song were reliably classified from acoustic features found across the 21 societies studied. The acoustic profiles of infant-directedness differed across language and music, but in a consistent fashion worldwide. In a secondary analysis, we studied whether listeners are sensitive to these acoustic features, playing the recordings to 51,065 people recruited online, from many countries, who guessed whether each vocalization was infant-directed. Their intuitions were largely accurate, predictable in part by acoustic features of the recordings, and robust to the effects of linguistic relatedness between vocalizer and listener. By uniting rich cross-cultural data with computational methods, we show links between the production of vocalizations and cross-species principles of bioacoustics, informing hypotheses of the psychological functions and evolution of human communication.

## Main

The forms of many animal signals are shaped by their functions, a link arising from production- and reception- related rules that help to maintain reliable signal detection within and across species^1–6^. Form-function links are widespread in vocal signals across taxa, from meerkats to fish^3, 7–10^, causing acoustic regularities that allow cross-species intelligibility^11–13, 15^. This facilitates the ability of some species to eavesdrop on the vocalizations of other species, for example, as in superb fairywrens (*Malurus cyaneus*), who learn to flee predatory birds in response to alarm calls that they themselves do not produce^14^.

In humans, an important context for the effective transmission of vocal signals is between parents and infants, as human infants are particularly helpless^16^. To elicit care, infants use a distinctive alarm signal: they cry^17^. In response, adults produce infant-directed language and music (sometimes called “parentese”) in forms of speech and song with putatively stereotyped acoustics^18–35^.

These stereotyped acoustics are thought to be functional: supporting language acquisition^36–39^, modulating infant affect and temperament^33, 40, 41^, and/or coordinating communicative interactions with infants^46–48^. These theories all share a key prediction: like the vocal signals of other species, the forms of infant-directed vocalizations should be shaped by their functions, instantiated with clear regularities across cultures. Put another way, we should expect people to *alter* the acoustics of their vocalizations when those vocalizations are directed toward infants, and they should make those alterations in similar fashions worldwide.

The evidentiary basis for such a claim is controversial, however, given the limited generalizability of individual ethnographic reports and laboratory studies^53^; small stimulus sets^54^; and a variety of counterexamples^55, 56, 58–62^. Some evidence suggests that infant-directed speech is primarily characterized by higher and more variable pitch^63^ and more exaggerated and variable vowels^23, 64, 65^, based on studies in modern industrialized societies^23, 28, 49, 50, 52, 66, 67^ and a few small-scale societies^51, 68^. Infants are themselves sensitive to these features, preferring them, even if spoken in unfamiliar languages^69–71^. But these acoustic features are less exaggerated or reportedly absent in some cultures^60, 66, 72^ and may vary in relation to the age and sex of the infant^66, 73, 74^, weighing against claims of cross-cultural regularities.

In music, infant-directed songs also seem to have some stereotyped acoustic features. Lullabies, for example, tend toward slower tempos, reduced accentuation, and simple repetitive melodic patterns^31, 32, 35, 75^, supporting functional roles associated with infant care^33, 41, 46^ in industrialized^34, 76–78^ and small-scale societies^79, 80^. Infants are soothed by these acoustic features, whether produced in familiar^44, 45^ or unfamiliar songs^81^, and both adults and children reliably associate the same features with a soothing function^31, 32, 75^. But cross-cultural studies of infant-directed song have primarily relied upon archival recordings from disparate sources^29, 31, 32^; an approach that poorly controls for differences in voices, behavioral contexts, recording equipment, and historical conventions, limiting the precision of findings and complicating their generalizability.

Measurements of the *same voices* producing multiple vocalizations, gathered from many people in many languages, worldwide, would enable the clearest analyses of whether and how humans alter the acoustics of their vocalizations when communicating with infants, helping to address the lack of consensus in the literature. Further, yoked analyses of both speech and song may explain how the forms of infant-directed vocalizations reliably differ from one another, testing theories of their shared or separate functions^33, 36–41, 46^.

We take this approach here. We built a corpus of infant-directed speech, adult-directed speech, infant-directed song, and adult-directed song from 21 human societies, totaling 1615 recordings of 410 voices (Fig. 1a, Table 1, and Methods; the corpus is open-access at https://doi.org/10.5281/zenodo.5525161). We aimed to maximize linguistic, cultural, geographic, and technological diversity: the recordings cover 18 languages from 11 language families and represent societies located on 6 continents, with varying degrees of isolation from global media, including 4 small-scale societies that lack access to television, radio, or the internet and therefore have strongly limited exposure to language and music from other societies. Participants were asked to provide all four vocalization types.

**Fig. 1.**
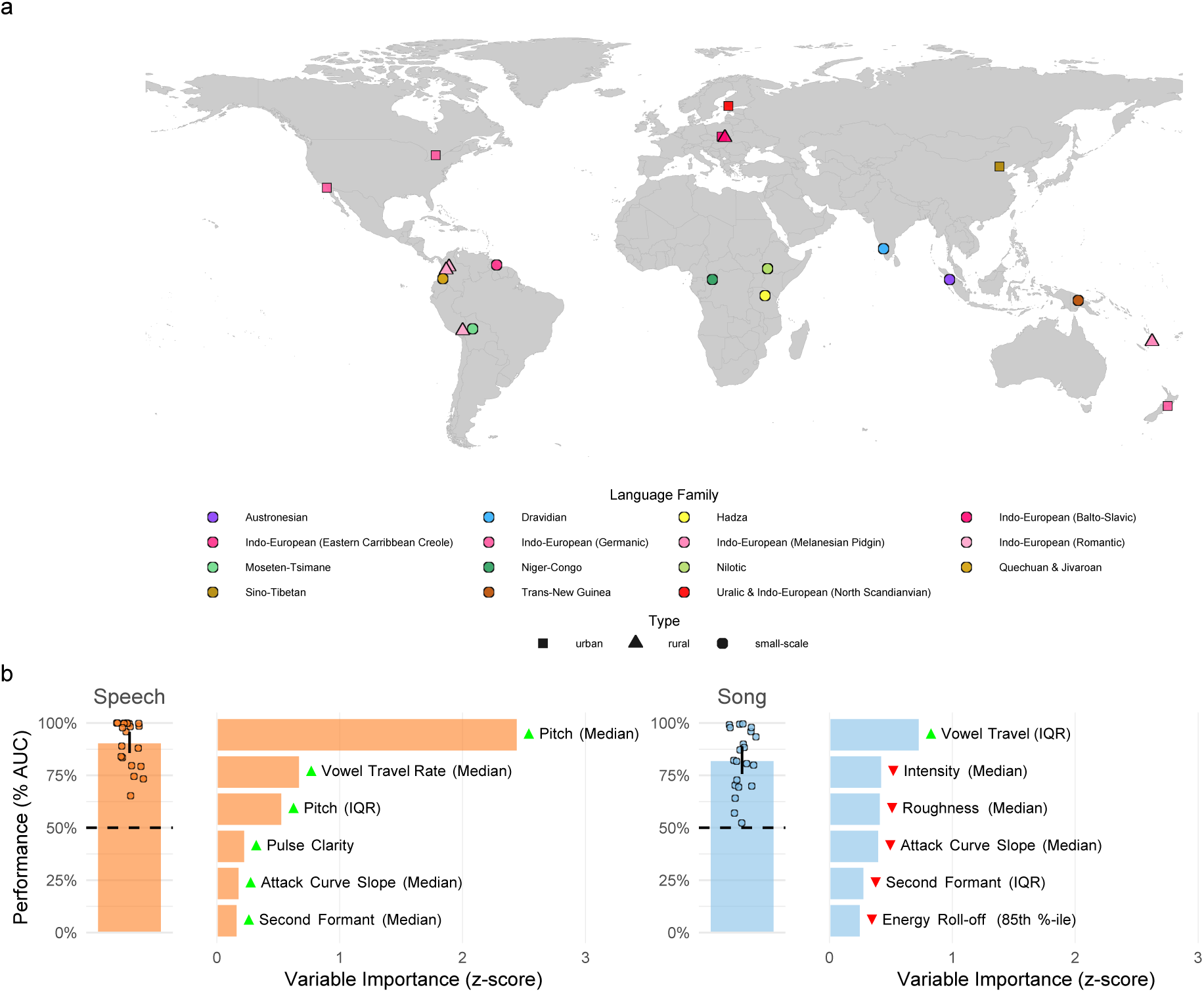
Cross-cultural regularities in infant-directed vocalizations. **a**, We recorded examples of speech and song from 21 urban, rural, or small-scale societies, in many languages. The map indicates the approximate locations of each society and is color-coded by the language family or sub-group represented by the society. **b**, Machine-learning classification demonstrates the stereotyped acoustics of infant-directed speech and song. We trained two least absolute shrinkage and selection operator (LASSO) models, one for speech and one for song, to classify whether recordings were infant-directed or adult-directed on the basis of their acoustic features. These predictors were regularized using fieldsite-wise cross-validation, such that the model optimally classified infant-directedness across all 21 societies studied. The vertical bars represent the overall classification performance (quantified via receiver operating characteristic/area under the curve; AUC); the error bars represent 95% confidence intervals; the points represent the performance estimate for each fieldsite; and the horizontal dashed lines indicate chance level of 50% AUC. The horizontal bars show the six acoustic features with the largest influence in each classifier; the green and red triangles indicate the direction of the effect, e.g., with median pitch having a large, positive effect on classification of infant-directed speech. The full results of the variable selection procedure are in Extended Data Table 2, with further details in Methods.

**Table 1.**
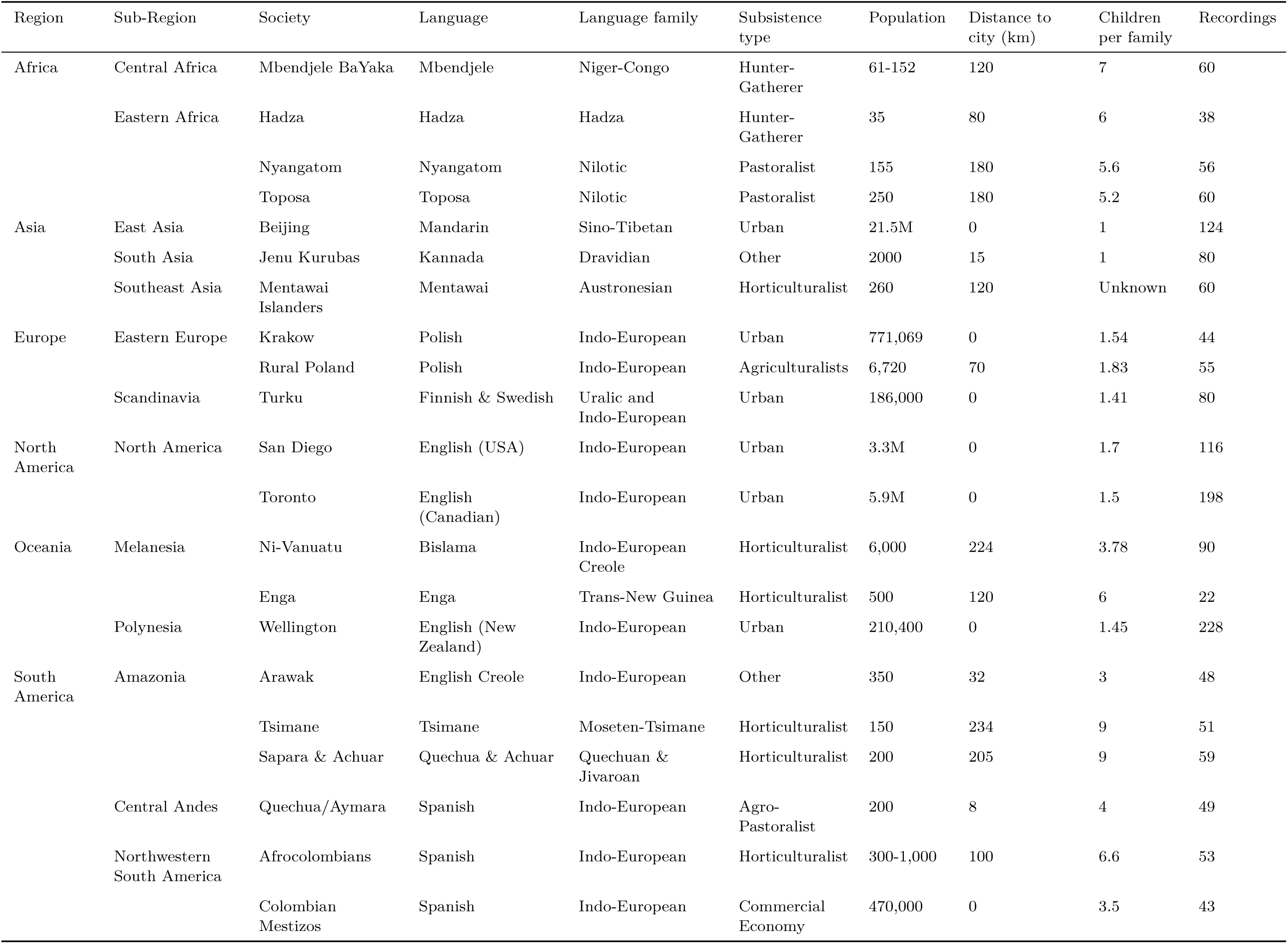
Societies from which recordings were gathered.

We used computational analyses of the acoustic forms of the vocalizations and a citizen-science experiment to test (i) the degree to which infant-directed vocalizations are cross-culturally stereotyped; and (ii) the degree to which naïve listeners detect infant-directedness in language and music.

## The acoustic forms of infant-directed speech and song are stereotyped across cultures

We studied 15 types of acoustic features in each recording (e.g., pitch, rhythm, timbre) via 94 summary variables (e.g., median, interquartile range) that were treated to reduce the influence of atypical observations (e.g., extreme values caused by loud wind, rain, and other background noises) (see Methods and SI Text 1.1; a codebook is in Extended Data Table 1). To minimize the potential for bias, we collected the acoustic data using automated signal extraction tools that measure physical characteristics of the auditory signal; such physical characteristics lack cultural information (in contrast to, e.g., human annotations) and thus can be applied reliably across diverse audio recordings.

First, we asked whether the acoustics of infant-directed speech and song are stereotyped in similar ways across the societies whose recordings we studied. Following previous work^32^, we used a least absolute shrinkage and selection operator (LASSO) logistic classifier^82^ with fieldsite-wise *k*-fold cross-validation, separately for speech and song recordings, using all 15 types of acoustic features (see Methods). This approach provides a strong test of cross-cultural regularity: the model is trained *only* on data from 20 of the 21 societies to predict whether each vocalization in the 21st society is infant- or adult-directed. The procedure is repeated 20 further times, with each society being held out, optimizing the model to maximize classification performance across the full set of societies. The summary of the model’s performance reflects, corpus-wide, the degree to which infant-directed speech and song are acoustically stereotyped, as high classification performance can only result from cross-cultural regularities.

The models accurately classified both speech and song, on average (Fig. 1b; speech: area under the curve, AUC = 91%, 95% CI [86%, 96%]; song: AUC = 82%, 95% CI [76%, 89%]). Evaluating classification performance within the recordings in each fieldsite showed a high degree of cross-cultural regularity, with the average performance in 21 of 21 fieldsites’ above chance level for both speech and song recordings (Fig. 1b).

To test the reliability of these findings, we repeated them with two alternate cross-validation strategies, using the same cross-validation procedure but doing so across language families and geographic regions instead of fieldsites. The results robustly replicated in both cases (Extended Data Fig. 1). Moreover, to ensure that the main LASSO results were not attributable to particulars of the audio-editing process (see Methods), we also repeated them using unedited audio from the corpus; the results replicated again (Extended Data Fig. 2).

These findings show that the acoustic features of infant-directed speech and song are robustly stereotyped across the 21 societies studied here.

### The acoustic profile of infant-directedness differs across speech and song

We used two convergent approaches to determine the specific acoustic features that are predictive of infant- directedness in speech and song.

First, the LASSO procedure identified the most reliable predictors of contrasts between infant- and adult-directed vocalizations. The most influential of these predictors are reported in Fig. 1b, with their relative variable importance scores, and show substantial differences in the variables the model relied upon to reliably classify speech and song across cultures. For example, pitch (F_0_ median and interquartile range) and median vowel travel rate strongly differentiated infant-directedness in speech, but not in song; while vowel travel variability (interquartile range) and median intensity strongly differentiated infant-directedness in song, but not in speech. The full results of the LASSO variable selection are in Extended Data Table 2.

Second, in a separate exploratory-confirmatory analysis, we used mixed-effects regression to measure the expected difference in each acoustic feature associated with infant-directedness, separately for speech and song. Importantly, this approach estimates main effects adjusted for sampling variability *and* estimates fieldsite-level effects, allowing for tests of the degree to which the main effects differ in magnitude across cultures (e.g., for a given acoustic feature, if recordings from some fieldsites show larger differences between infant- and adult-directed speech than do recordings from other fieldsites). The analysis was preregistered.

The procedure identified 11 acoustic features that reliably distinguished infant-directedness in song, speech, or both (Fig. 2; statistics are in Extended Data Table 3); we also estimated these effects within each fieldsite (see the doughnut plots in Fig. 2 and full estimates in Extended Data Table 4).

**Fig. 2.**
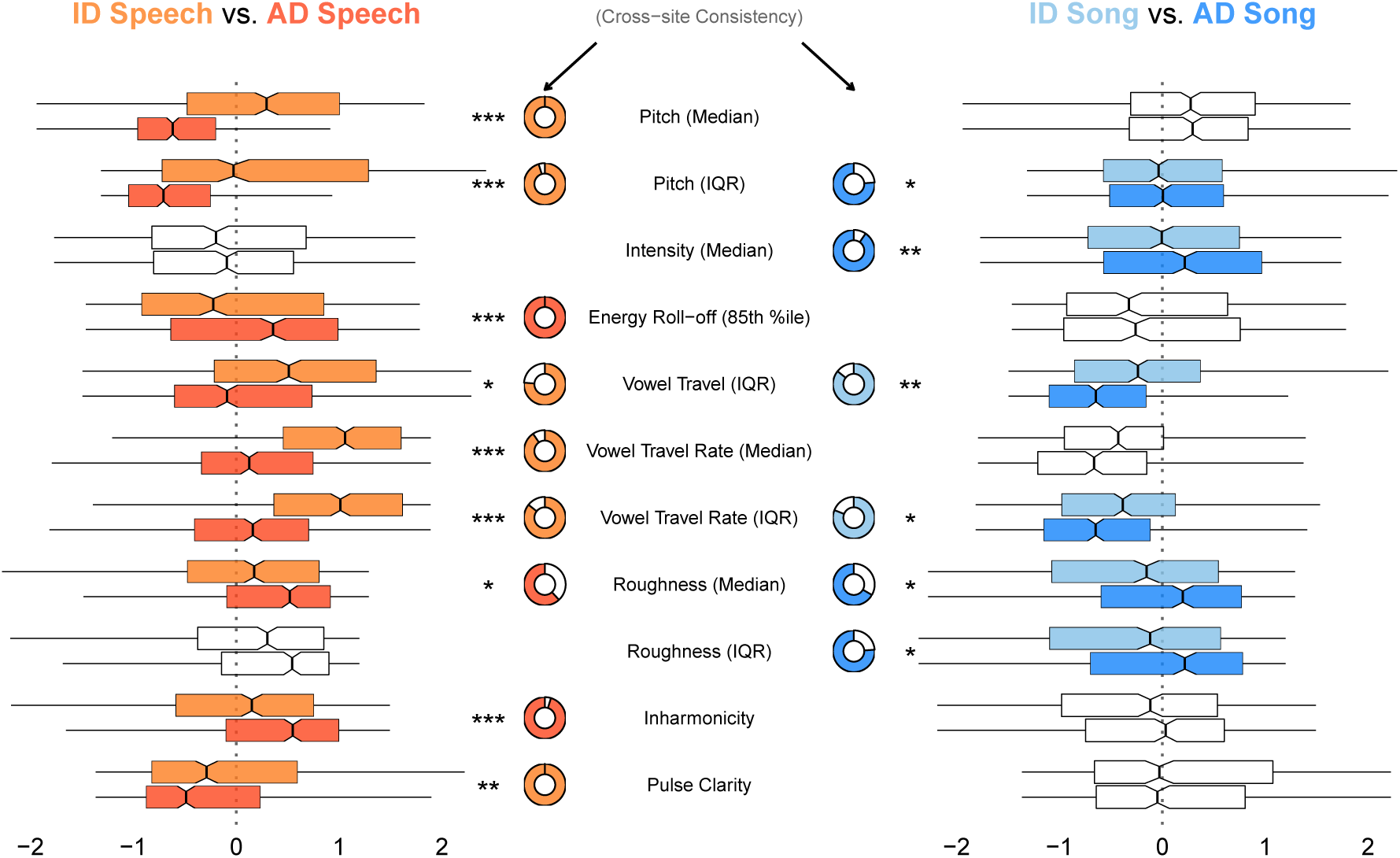
How people alter their voices when vocalizing to infants. Eleven acoustic features had a statistically significant difference between infant-directed and adult-directed vocalizations, within-voices, in speech, song, or both. Consistent with the LASSO results (Fig. 1b and Extended Data Table 2), the acoustic features operated differently across speech and song. For example, median pitch was far higher in infant-directed speech than in adult-directed speech, whereas median pitch was comparable across both forms of song. Some features were highly consistent across fieldsites (e.g., lower inharmonicity in infant-directed speech than adult-directed speech), whereas others were more variable (e.g., lower roughness in infant-directed speech than adult-directed speech). The boxplots, which are ordered approximately from largest to smallest differences between effects across speech and song, represent each acoustic feature’s median (vertical black lines) and interquartile range (boxes); the whiskers indicate 1.5 *×* IQR; the notches represent the 95% confidence intervals of the medians; and the doughnut plots represent the proportion of fieldsites where the main effect repeated, based on estimates of fieldsite-wise random effects. Only comparisons that survived an exploratory-confirmatory analysis procedure are plotted; faded comparisons did not reach significance in confirmatory analyses. Significance values are computed via linear combinations, following multi-level mixed-effects models; \**p <* 0.05, \*\**p <* 0.01, \*\*\**p <* 0.001. Regression results are in Extended Data Table 3 and full reporting of fieldsite-level estimates is in Extended Data Table 4. Note: the model estimates are normalized jointly on speech and song data so as to enable comparisons *across* speech and song for each feature; as such, the absolute distance from 0 for a given feature is not directly interpretable, but estimates are directly comparable across speech and song.

In speech, across all or the majority of societies, infant-directedness was characterized by higher pitch, greater pitch range, and more contrasting vowels than adult-directed speech from the same voices (largely replicating the results of the LASSO approach; Fig. 1b and Extended Data Table 2). Several acoustic effects were consistent in all fieldsites (e.g., pitch, energy roll-off, pulse clarity), while other features, such as vowel contrasts and inharmonicity were consistent in the majority of them. These patterns align with prior claims of pitch and vowel-contrast being robust features of infant-directed speech^23, 67^, and substantiate them across many cultures.

The distinguishing features of infant-directed song were more subtle than those of speech but nevertheless corroborate its purported soothing functions^33, 41, 46^: reduced intensity and acoustic roughness, although these were less consistent across fieldsites than the speech results. The less-consistent effects may result from the fact that while solo-voice speaking is fairly natural and representative of most adult-directed speech (i.e., people rarely speak at the same time), much of the world’s song occurs in social groups where there are multiple singers and accompanying instruments^32, 46, 83^. Asking participants to produce solo adult-directed song may have biased participants toward choosing more soothing and intimate songs (e.g., ballads, love songs; see Extended Data Table 5) or less naturalistic renditions of songs; and the production of songs in the presence of an infant, which could potentially alter participants’ singing style^35^. Thus, the distinctiveness of infant-directed song (relative to adult-directed song) may be underestimated here.

The exploratory-confirmatory analyses provided convergent evidence for opposing acoustic trends across infant-directed speech and song, as did an alternate approach using principal-components analysis; three principal components most strongly distinguished speech from song, infant-directed song from adult-directed song, and infant-directed speech from adult-directed speech (SI Text 1.2 and Extended Data Fig. 3). Replicating the LASSO findings, for example, median pitch strongly differentiated infant-directed speech from adult-directed speech, but it had no such effect in music; pitch variability had the *opposite* effect across language and music; and further differences were evident in pulse clarity, inharmonicity, and energy roll-off. These patterns are consistent with the possibility of differentiated functional roles across infant-directed speech and song^18, 33, 34, 45, 46, 81, 84^.

Some acoustic features were nevertheless common to both language and music; in particular, overall, infant- directedness was characterized by reduced roughness, which may facilitate parent-infant signalling^5, 41^ through better contrast with the sounds of screaming and crying^17, 85^; and increased vowel contrasts, potentially to aid language acquisition^36, 37, 39^ or as a byproduct of socio-emotional signalling^1, 65^.

### Naïve listeners are sensitive to the acoustic forms of infant-directed vocalizations

If people worldwide reliably alter their speech and song when interacting with infants, as the above findings demonstrate, this may enable listeners to make reliable inferences concerning the intended targets of speech and song, consistent with functional accounts of infant-directed vocalization^33, 36–43, 46^. We tested this secondary hypothesis in a simple listening experiment, conducted in English using web-based citizen-science methods^86^.

We played excerpts from the vocalization corpus to 51,065 people in the “Who’s Listening?” game on https://themusiclab.org (after exclusions; see Methods). The participants resided in 187 countries (Fig. 3b) and reported speaking 199 languages fluently (including second languages, for bilinguals). We asked them to judge, quickly, whether each vocalization was directed to a baby or to an adult (see Methods and Extended Data Fig. 4).

**Fig. 3.**
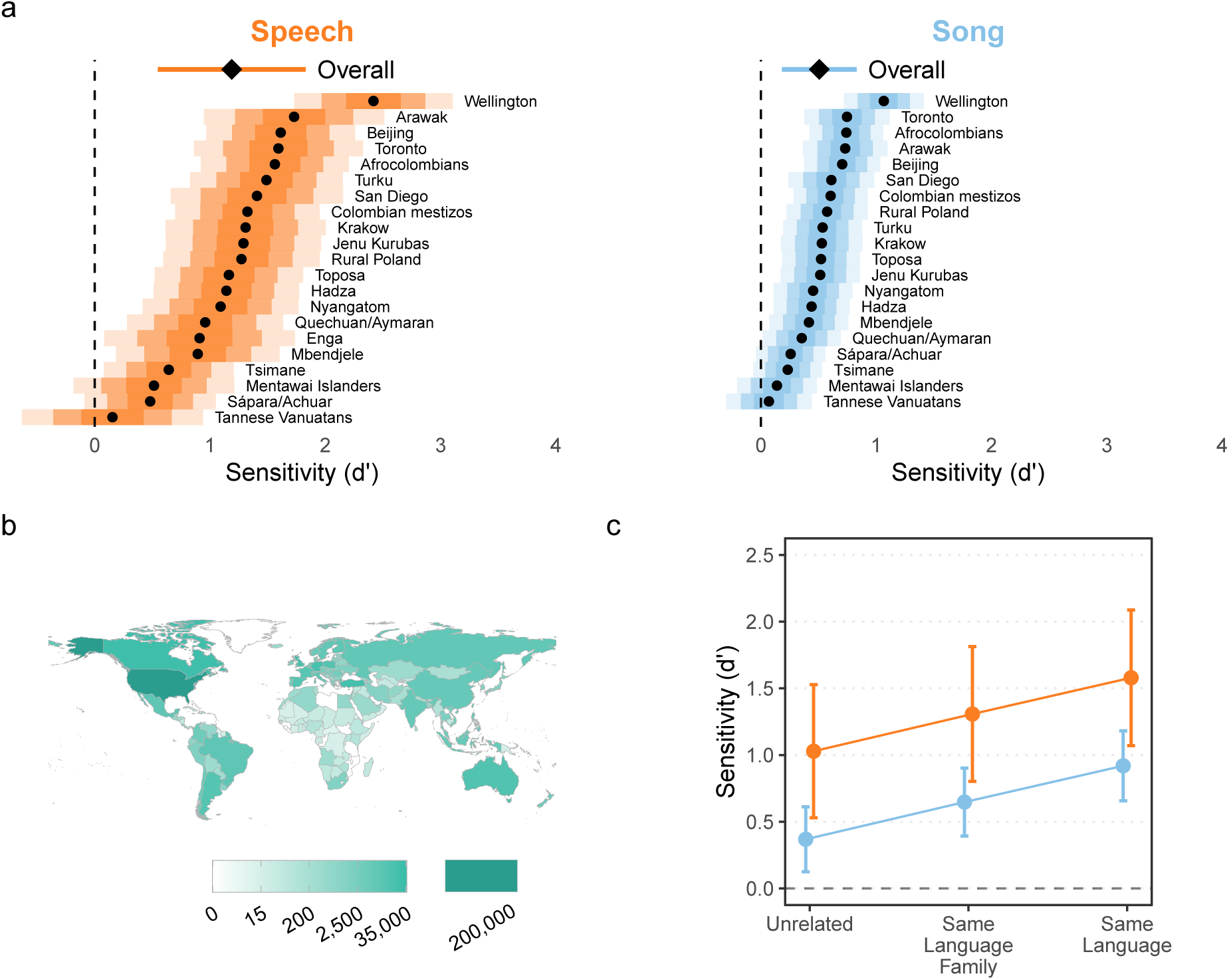
Naïve listeners distinguish infant-directed vocalizations from adult-directed vocalizations across cultures. Participants listened to vocalizations drawn at random from the corpus, viewing the prompt “Someone is speaking or singing. Who do you think they are singing or speaking to?” They could respond with either “adult” or “baby” (Extended Data Fig. 4). From these ratings, we computed listener sensitivity (*d^′^*). **a**, Listeners reliably detected infant-directedness in both speech and song, overall (indicated by the diamonds, with 95% confidence intervals indicated by the horizontal lines), and across many fieldsites (indicated by the black dots), although the strength of the fieldsite-wise effects varied substantially (see the distance between the vertical dashed line and the black dots; the shaded regions represent 50%, 80%, and 95% confidence intervals, in increasing order of lightness). Note that one fieldsite-wise *d^′^* could not be estimated for song; complete statistical reporting is in Extended Data Table 6. **b**, The participants in the citizen-science experiment hailed from many countries; the gradients indicate the total number of vocalization ratings gathered from each country. **c**, The main effects held across different combinations of the linguistic backgrounds of vocalizer and listener. We split all trials from the main experiment into three groups: those where a language the listener spoke fluently was the same as the language of the vocalization (*n* = 82,094; those where a language the listener spoke fluently was in the same major language family as the language of the vocalization (*n* = 110,664), and those with neither type of relation (*n* = 285,378). The plot shows the estimated marginal effects of a mixed-effects model predicting *d^′^* values across language and music examples, after adjusting for fieldsite-level effects. In all three cases, the main effects replicated; increases in linguistic relatedness corresponded with increases in sensitivity.

The responses were strongly biased toward “baby” responses when hearing songs and away from “baby” responses when hearing speech, regardless of the actual target of the vocalizations (Extended Data Fig. 5). To correct for these response biases, we used *d*-prime analyses at the level of each vocalist, i.e., analyzing listeners’ sensitivity to infant-directedness in speech and song (SI Text 1.3). Unless noted otherwise, all estimates reported here are generated by mixed-effects linear regression, adjusting for fieldsite nested within world region, via random effects.

The listeners’ intuitions were accurate, on average and across fieldsites (Fig. 3a). Sensitivity (*d^′^*) was significantly higher than the chance level of 0 (speech: *d^′^* = 1.19, 95% CI [0.55, 1.83]; song: *d^′^* = 0.51, 95% CI [0.18, 0.83]; *ps < .*05). These results were robust to learning effects (Extended Data Fig. 6) and to multiple data trimming decisions. For example, they replicated whether or not recordings with confounding contextual/background cues (e.g., an audible infant) were excluded and also when data from English-language recordings, which might be understandable to participants, were excluded (SI Text 1.4).

To test the consistency of listener inferences across cultures, we estimated fieldsite-level sensitivity from the random effects in the model. Cross-site variability was evident in the magnitude of sensitivity effects: listeners were far better at detecting infant-directedness in some sites than others (with very high *d^′^* in the Wellington, New Zealand site for both speech and song, but marginal *d^′^* in Tannese Vanuatans, for example). Nevertheless, the estimated mean fieldsite-wise *d^′^* was greater than 0 in both speech and song in all fieldsites (Fig. 3a); with 95% confidence intervals not overlapping with 0 in 18 of 21 fieldsites for speech and 16 of 20 for song (Extended Data Table 6; one *d^′^* estimate could not be computed for song due to missing data). Most fieldsite-wise sample sizes after exclusions were small (see Methods), so we caution that fieldsite-wise estimates are far less interpretable than the overall *d^′^* estimate reported above.

Analyses of cross-cultural variability among *listeners* revealed similarities in their perception of infant- directedness. In particular, coefficient of variation scores revealed little variation in listener accuracy across countries of origin (2.3%) and native languages (1.1%), with the estimated effects of age and gender both less than 1%. And more detailed demographic characteristics available for a subset of participants in the United States, including socioeconomic status and ethnicity, also explained little variation in accuracy (SI Text 1.5). These findings suggest general cross-demographic consistency in listener intuitions.

One important aspect of listeners was predictive of their performance, however: their degree of relatedness to the vocalizer, on a given trial. To analyze this, we estimated fixed effects for three forms of linguistic relatedness between listener and vocalizer: (i) *weak relatedness*, when a language the listener spoke fluently was from a different language family than that of the vocalization (e.g., when the vocalization was in Mentawai, an Austronesian language, and the listener’s native language was Mandarin, a Sino-Tibetan language); (ii) *moderate relatedness*, when the languages were from the same language family (e.g., when the vocalization was in Spanish and the listener spoke fluent English, which are both Indo-European languages); or (iii) *strong relatedness*, when a language the listener spoke fluently exactly matched the language of the vocalization.

Sensitivity was significantly above chance in all cases (Fig. 3c), with increases in performance associated with increasing relatedness (unrelated: estimated speech *d^′^* = 1.03, song *d^′^* = 0.37; same language family: speech *d^′^* = 1.31, song *d^′^* = 0.65; same language: speech *d^′^* = 1.58, song *d^′^* = 0.92). Some of this variability is likely attributable to trivial language comprehensiblity (i.e., in cases of strong relatedness, listeners very likely understood the words of the vocalization, strongly shaping their infant-directedness rating).

These findings provide an important control, as they demonstrate that the overall effects (Fig. 3a) are not attributable to linguistic similarities between listeners and vocalizers (Fig. 3c), which could, for example, allow listeners to detect infant-directedness on the basis of the words or other linguistic features of the vocalizations, as opposed to their *acoustic* features. And while the experiment’s instructions were presented in English (suggesting that all listeners likely had at least a cursory understanding of English), the findings were robust to the exclusion of all English-language recordings (SI Text 1.4).

We also found suggestive evidence of other, non-linguistic links between listeners and vocalizers being predictive of sensitivity. For example, fieldsite population size and distance to the nearest urban center were correlated estimated sensitivity to infant-directedness in that fieldsite. These and similar effects (reported in SI Text 1.6) suggests that performance was somewhat higher in the larger, more industrialized fieldsites that are more similar to the environments of internet users, on average. But these analyses are necessarily coarser than the linguistic relatedness tests reported above.

### Human intuitions of infant-directedness are modulated by vocalization acoustics

Last, we studied the degree to which the acoustic features of the recordings were predictive of listeners’ intuitions concerning them (measured as the experiment-wide *proportions* of infant-directedness ratings for each vocalization, in a similar approach to other research^75^). These proportions can be considered a continuous measure of perceived infant-directedness, per the ears of the naïve listeners. We trained two LASSO models to predict the proportions, with the same fieldsite-wise cross-validation procedure used in the acoustic analyses reported above. Both models explained variation in human listeners’ intuitions, albeit more so in speech than in song (Fig. 4; speech *R*^2^ = 0.59; song *R*^2^ = 0.18, *ps* < 0.0001), likely because the acoustic features studied here more weakly guided listeners’ intuitions in song than they did in speech.

**Fig. 4.**
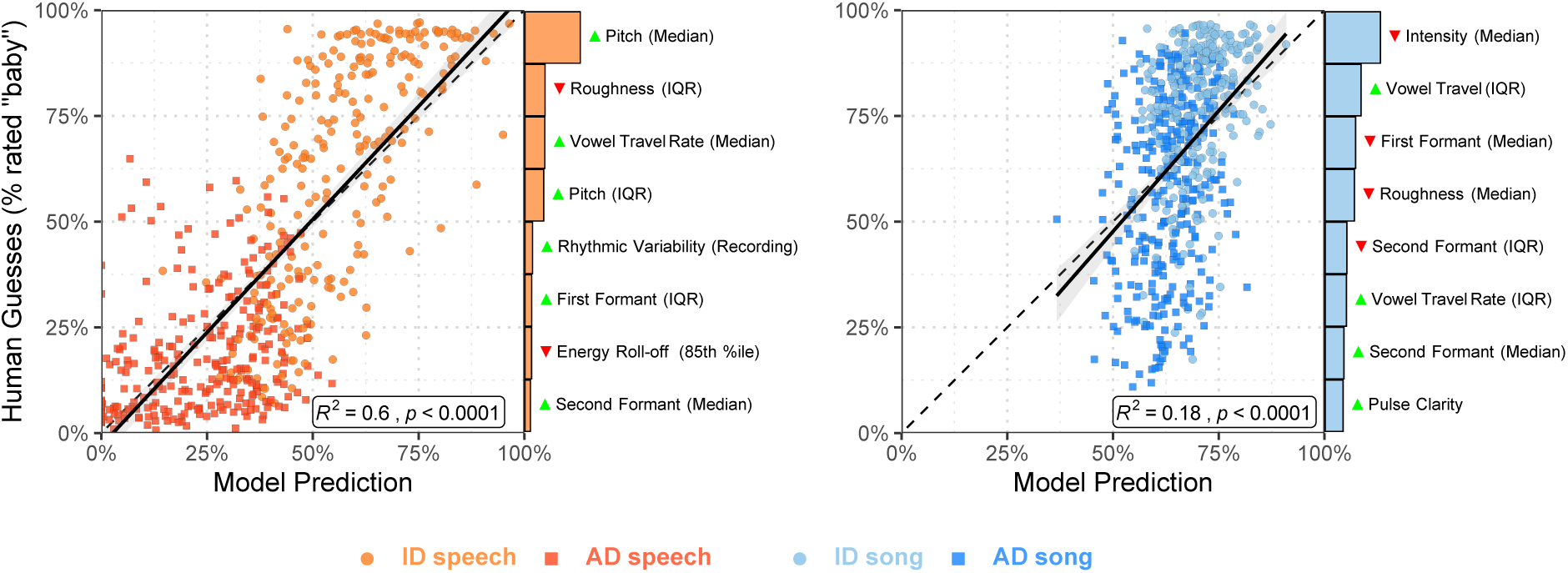
Human inferences about infant-directedness are predictable from acoustic features of vocalizations. To examine the degree to which human inferences were linked to the acoustic forms of the vocalizations, we trained two LASSO models to predict the proportion of “baby” responses for each recording from the human listeners. While both models explained substantial variability in human responses, the model for speech was more accurate than the model for song, in part because the human listeners erroneously relied on acoustic features for their predictions in song that less reliably characterized infant-directed song across cultures (see Figs. 1b and 2). Each point represents a recorded vocalization, plotted in terms of the model’s estimated infant-directedness of the model and the average “infant-directed” rating from the naïve listeners; the barplots depict the relative explanatory power of the top 8 acoustical features in each LASSO model, showing which features were most strongly associated with human inferences (the green or red triangles indicate the directions of effects, with green higher in infant-directed vocalizations and red lower); the dotted diagonal lines represent a hypothetical perfect match between model predictions and human guesses; the solid black lines depict linear regressions; and the grey ribbons represent the standard errors of the mean, from the regressions.

If human inferences are attuned to cross-culturally reliable acoustic correlates of infant-directedness, one might expect a close relationship between the strength of *actual* acoustic differences between vocalizations on a given feature and the relative influence of that feature on human intuitions. To test this question, we correlated how strongly a given acoustic feature distinguished infant-directed from adult-directed speech and song (Fig. 2; estimated with mixed-effects modeling) with the variable importance of that feature in the LASSO model trained to predict human intuitions (the bar plots in Fig. 4). We found a significant strong positive relationship for speech (*r* = 0.72, *p* = 0.001) but not for song (*r* = 0.36, *p* = 0.08).

This difference may help to explain the weaker intuitions of the naïve listeners in song, relative to speech: naïve listeners’ inferences about speech were more directly driven by acoustic features that *actually* characterize infant-directed speech worldwide, whereas their inferences about song were erroneously driven by acoustic features that *less reliably* characterize infant-directed song worldwide. For example, songs with higher pulse clarity and median second formats, and lower median first formants were more likely to be rated as infant-directed, but these features did not reliably correlate with infant-directed song across cultures in the corpus (and, accordingly, neither approach to the acoustic analyses identified them as reliable correlates of infant-directedness in music). Intuitions concerning infant-directed song may also have been driven by more subjective features of the recordings, higher-level acoustic features that we did not measure, or both.

We note, however, that the interpretation of this difference may be limited by the representativeness of the sample of recordings: the differences in the models’ ability to predict listeners’ intuitions could alternatively be driven by differences in the true representativeness of one or more of the vocalization types.

## Discussion

We provide convergent evidence for cross-cultural regularities in the acoustic design of infant-directed speech and song. Infant-directedness was robustly characterized by core sets of acoustic features, across the 21 societies studied, and these sets of features differed reliably across speech and song. Naïve listeners were sensitive to the acoustical regularities, as they reliably identified infant-directed vocalizations as more infant-directed than adult-directed vocalizations, despite the fact that the vocalizations were of largely unfamiliar cultural, geographic, and linguistic origin.

Thus, despite evident variability in language, music, and infant care practices worldwide, when people speak or sing to fussy infants, they modify the acoustic features of their vocalizations in similar and mutually intelligible ways across cultures. This evidence supports the hypothesis that the forms of infant-directed vocalizations are shaped by their functions, in a fashion similar to the vocal signals of many non-human species.

These findings do not mean that infant-directed speech and song always sound the same across cultures. Indeed, the classification accuracy of a machine-learning model varied, with some fieldsites demonstrating larger acoustic differences between infant- and adult-directed vocalizations than other fieldsites. Similarly, the citizen-science participants’ ratings of infant-directedness differed substantially in magnitude across fieldsites. But such variability also does not imply the *absence* of cross-cultural regularities. Instead, they support an account of acoustic variation stemming from epigenetic rules: species-typical traits which bias cultural variation in one direction rather than another^87^. Put another way, the pattern of evidence strongly implies a core set of cross-cultural acoustic and perceptual regularities which are also shaped by culture.

By analyzing both speech and song recorded from the same voices, we discerned precise differences in the ways infant-directedness is instantiated in language and music. In response to the same prompt of addressing a “fussy infant”, infant-directedness in speech and song was instantiated with opposite trends in acoustic modification (relative to adult-directed speech and song, respectively): infant-directed speech was more intense and contrasting (e.g., more pitch variability, higher intensity) while infant-directed song was more subdued and soothing (e.g., less pitch variability, lower intensity). These acoustic dissociations comport with functional dissociations, with speech being more attention-grabbing, the better to distract from baby’s fussiness^37, 38^; and song being more soothing, the better to lower baby’s arousal^32, 33, 41–43, 45, 81^. Speech and song are both capable of playful or soothing roles^62^ but each here tended toward one acoustic profile over the other, despite both types of vocalization being elicited here in the *same* context: vocalizations used “when the baby is fussy”.

Many of the reported acoustic differences are consistent with properties of vocal signalling in non-human animals, raising the intriguing possibility that the designs of human communication systems are rooted in the basic principles of bioacoustics^1–15^. For example, in both speech and song, infant-directedness was robustly associated with purer and less harsh vocal timbres, and greater formant-frequency dispersion (expanded vowel space). And in speech, one of the largest and most cross-culturally robust effects of infant-directedness was higher pitch (F_0_). In non-human animals, these features have convergently evolved across taxa in the functional context of signalling friendliness or approachability in close contact calls^1, 3, 65, 88^, in contrast to alarm calls or signals of aggression, which are associated with low-pitched, rough sounds with less formant dispersal^4, 89–91^. The use of these features in infant care may originate from signalling approachability to baby, but may have later acquired further functions more specific to the human developmental context. For example, greater formant-frequency dispersion accentuates vowel contrasts, which could facilitate language acquisition^36, 65, 92–94^; and purer vocal timbre may facilitate communication by contrasting conspicuously with the acoustic context of infant cries^5^ (for readers unfamiliar with infants, their cries are acoustically harsh^17, 85^).

Such conspicuous contrasts may have the effect of altering speech to make it more song-like when interacting with infants, as Fernald^18^ notes: *“. . . the communicative force of [parental] vocalizations derive not from their arbitrary meanings in a linguistic code, but more from their immediate musical power to arouse and alert, to calm, and to delight”*.

Comparisons of the acoustic effects across speech and song reported here support this idea. Infant-directedness altered the pitch level (F_0_) of speech, bringing it roughly to a level typical of song, while also increasing pulse clarity. These characteristics of music have been argued to originate from elaborations to infant-directed vocalizations, where both use less harsh but more variable pitch patterns, more temporally variable and expansive vowel spaces, and attention-orienting rhythmic cues to provide infants with ostensible “flashy” signals of attention and pro-social friendliness^41, 46, 63, 95, 96^. Pitch alterations are not *absent* from infant-directed song, of course; in one study, mothers sang a song at higher pitch when producing a more playful rendition, and a lower pitch when producing a more soothing rendition^44^. But on average, both infant- and adult-directed song, along with infant-directed speech, tend to be higher in pitch than adult-directed speech. In sum: the constellation of acoustic features that characterize infant-directedness in speech, across cultures, are rather musical.

We leave open at least four sets of further questions. First, the results are suggestive of universality in the production of infant-directed vocalizations, because the corpus covers a swath of geographic locations (21 societies on 6 continents), languages (12 language families), and different subsistence regimes (8 types) (see Table 1). But the participants studied do not constitute a representative sample of humans (nor do the societies or languages studied constitute a representative sample of human societies or languages), so a strong claim of universality would not be justified. Future work may assess the validity of such a universality claim by studying infant-directed vocalizations in a wider range of human societies, and by using phylogenetic methods to examine whether people in societies that are distantly related nonetheless produce similar infant-directed vocalizations.

Second, the naïve listener experiment tested orders of magnitude more participants and covering a far more diverse set of countries and native languages than did prior research, raising the possibility that they may generalize across many populations. But such a generalization is not fully justified, because the instructions of the experiment were presented in English, on an English-language website. Future work may determine their generality by testing perceived infant-directedness in multilingual experiments, to more accurately characterize cross-cultural variability in the perception of infant-directedness; and by testing listener intuitions among groups with reduced exposure to a given set of infant-directed vocalizations, such as very young infants or people from isolated, distantly related societies, as in related efforts^27, 69, 97^. Such research would benefit in particular from a focus on societies previously reported to have unusual vocalization practices, infant care practices, or both^55, 58–60^; and would also clarify the extent to which convergent practices across cultures are due to cultural borrowing (in the many cases where societies are not fully isolated from the influence of global media).

Third, most prior studies of infant-directed vocalizations use *elicited* recordings^20, 23, 26, 30, 39, 44^, as did we. While this method may underestimate the differences between infant-directed and adult-directed vocalizations, whether and how simulated infant-directed speech and song differ from their naturalistic counterparts is poorly understood. Future work may explore this issue by analyzing recordings of infant-directed vocalizations that are covertly and/or unobtrusively collected in a non-elicited manner, as in research using wearable recording devices for infants^78, 98^. This may also resolve potential confounds from the wording of instructions to vocalizers.

Last, we note that speech and song are used in multiple contexts with infants, of which “addressing a fussy infant” is just one^18, 34^. One curious finding may bear on general questions of the psychological functions of music: naïve listeners displayed a bias toward “adult” guesses for speech and “baby” guesses for song, regardless of their actual targets. We speculate that listeners treated “adult” and “baby” as the default reference levels for speech and song, respectively, against which acoustic evidence was compared, a pattern consistent with theories that posit song as having a special connection to infant care in human psychology^33, 46^.

## Methods

### Vocalization corpus

We built a corpus of 1,615 recordings of infant-directed song, infant-directed speech, adult-directed song, and adult-directed speech (all audio is available at https://doi.org/10.5281/zenodo.5525161). Participants (*N* = 411) living in 21 societies (Fig. 1a and Table 1) produced each of these vocalizations, respectively, with a median of 15 participants per society (range 6-57). From those participants for whom information was available, most were female (86%) and nearly all were parents or grandparents of the focal infant (95%). Audio for one or more examples was unavailable from a small minority of participants, in cases of equipment failure or when the participant declined to complete the full recording session (25 recordings, or 1.5% of the corpus, were missing).

Recordings were collected by principal investigators and/or staff at their field sites, all using the same data collection protocol. They translated instructions to the native language of the participants, following the standard research practices at each site. There was no procedure for screening out participants, but we encouraged our collaborators to collect data from parents rather than non-parents. Fieldsites were selected partly by convenience (i.e., via recruiting principal investigators at fieldsites with access to infants and caregivers) and partly to maximize cultural, linguistic, and geographic diversity (see Table 1).

For infant-directed song and infant-directed speech, participants were asked to sing and speak to their infant as if they were fussy, where “fussy” could refer to anything from frowning or mild whimpering to a full tantrum. At no fieldsites were difficulties reported in the translation of the English word “fussy”, suggesting that participants understood it. For adult-directed speech, participants spoke to the researcher about a topic of their choice (e.g., they described their daily routine). For adult-directed song, participants sang a song that was not intended for infants; they also stated what that song was intended for (e.g., “a celebration song”). Participants vocalized in the primary language of their fieldsite, with a few exceptions (e.g., when singing songs without words; or in locations that used multiple languages, such as Turku, which included both Finnish and Swedish speakers).

For most participants (90%) an infant was physically present during the recording (the infants were 48% female; age in months: *M* = 11.40; SD = 7.61; range 0.5-48). When an infant was not present, participants were asked to imagine that they were vocalizing to their own infant or grandchild, and simulated their infant-directed vocalizations (a brief discussion is in SI Text 1.7).

In all cases, participants were free to determine the content of their vocalizations. This was intentional: imposing a specific content category on their vocalizations (e.g., “sing a *lullaby*”) would likely alter the acoustic features of their vocalizations, which are known to be influenced by experimental contexts^99^. Some participants produced adult-directed songs that shared features with the intended soothing nature of the infant-directed songs; data on the intended behavioral context of each adult-directed song are in Extended Data Table 5.

All recordings were made with Zoom H2n digital audio recorders, using foam windscreens (where available). To ensure that participants were audible along with researchers, who stated information about the participant and environment before and after the vocalizations, recordings were made with a 360° dual *x-y* microphone pattern. This produced two uncompressed stereo audio files (WAV) per participant at 44.1 kHz; we only analyzed audio from the two-channel file on which the participant was loudest.

The principal investigator at each fieldsite provided standardized background data on the behavior and cultural practices of the society (e.g., whether there was access to mobile-phones/TV/radio, and how commonly people used ID speech or song in their daily lives). Most items were based on variables included in the D-PLACE cross-cultural corpus^100^.

The 21 societies varied widely in their characteristics, from cities with millions of residents (Beijing) to small-scale hunter-gatherer groups of as few as 35 people (Hadza). All of the small-scale societies studied had limited access to TV, radio, and the internet, mitigating against the influence of exposure to the music and/or infant care practices of other societies. Four of the small-scale societies (Nyangatom, Toposa, Sápara/Achuar, and Mbendjele) were completely without access to these communication technologies.

The societies also varied in the prevalence of infant-directed speech and song in day-to-day life. The only site reported to lack infant-directed song in contemporary practice was the Quechuan/Aymaran site, although it was also noted that people from this site know infant-directed songs in Spanish and use other vocalizations to calm infants. Conversely, the Mbendjele BaYaka were noted to use infant-directed song, but rarely used infant-directed speech. In most sites, the frequency of infant-directed song and speech varied. For example, among the Tsimane, song was reportedly infrequent in the context of infant care; when it appears, however, it is apparently used to soothe and encourage infants to sleep.

Our default strategy was to analyze all available audio from the corpus. In some cases, however, this was inadvisable (e.g., in the naïve listener experiment, when a listener might understand the language of the recording, and make a judgment based on the recording’s linguistic content rather than its acoustic content); all exclusion decisions are explicitly stated throughout.

### Acoustic analyses

#### Acoustic feature extraction

We manually extracted the longest continuous and uninterrupted section of audio from each recording (i.e., isolating vocalizations by the participant from interruptions from other speakers, the infant, and so on), using Adobe Audition. We then used the silence detection tool in Praat^101^, with minimum sounding intervals at 0.1 seconds and minimum silent intervals at 0.3 seconds, to remove all portions of the audio where the participant was not speaking (i.e., the silence between vocalization phrases). These were manually concatenated in Python, producing denoised recordings, which were subsequently checked manually to ensure minimal loss of content.

We extracted and subsequently analyzed acoustic features using Praat^101^, MIRtoolbox^102^, temporal modularity using discrete Fourier transforms for rhythmic variability^103^, and normalized pairwise variability indices^104^. These features consisted of measurements of pitch (e.g., F_0_, the fundamental frequency), timbre (e.g., roughness), and rhythm (e.g., tempo; n.b., because temporal measures would be affected by the concatenation process, we computed these variables on unconcatenated audio only); all summarized over time: producing 94 variables in total. We standardized feature values within-voices, eliminating between-voice variability. Further technical details are in SI Text 1.1.

For both the LASSO analyses (Fig. 1b) and the regression-based acoustic analyses (Fig. 2), we restricted the variable set to 27 summary statistics of median and interquartile range, as these correlated highly with other summary statistics (e.g., maximum, range) but were less sensitive to extreme observations.

#### LASSO modeling

We trained least absolute shrinkage and selection operator (LASSO) logistic classifiers with cross-validation using tidymodels^105^. For both speech and song, these models were provided with the set of 27 acoustic variables described in the previous section. These raw features were then demeaned for speech and song separately within-voices and then normalized at the level of the whole corpus. During model training, multinomial log-loss was used as an evaluation metric to fit the lambda parameter of the model.

For the main analyses (Fig. 1b, Extended Data Table 2, and Extended Data Fig. 2) we used a *k*-fold cross-validation procedure at the level of fieldsites. Alternate approaches used *k*-fold cross-validation at the levels of language family and world region (Extended Data Fig. 1). We evaluated model performance using a receiver operating characteristic metric, binary area-under-the-curve (AUC). This metric is commonly used to evaluate the diagnostic ability of a binary classifier; it yields a score between 0% and 100%, with a chance level of 50%.

#### Mixed-effects modeling

Following a preregistered exploratory-confirmatory design, we fitted a multi-level mixed-effects regression predicting each acoustic variable from the vocalization types, after adjusting for voice and fieldsite as random effects, and allowing them to vary for each vocalization type separately. To reduce the risk of Type I error, we performed this analysis on a randomly selected half of the corpus (exploratory, weighting by fieldsite) and only report results that successfully replicated in the other half (confirmatory). We did not correct for multiple tests because the exploratory-confirmatory design restricts the tests to those with a directional prediction.

These analyses deviated from the preregistration in two minor ways. First, we retained planned comparisons within vocalization types, but we eliminated those that compared across speech and song when we found much larger acoustic differences between speech and song overall than the differences between infant- and adult-directed vocalizations (a fact we failed to predict). As such, we adopted the simpler approach of post-hoc comparisons that were only within speech and within song. For transparency, we still report the preregistered post-hoc tests in Extended Data Fig. 7, but suggest that these comparisons be interpreted with caution. Second, to enable fieldsite-wise estimates (reported in Extended Data Table 4), we normalized the acoustic data corpus-wide and included a random effect of participant, rather than normalizing within-voices (as within-voice normalization would set all fieldsite-level effects to 0, making cross-fieldsite comparisons impossible).

### Naïve listener experiment

We analyzed all data available at the time of writing this paper from the “Who’s Listening?” game at https://themusiclab.org/quizzes/ids, a continuously running jsPsych^106^ experiment distributed via Pushkin^107^, a platform that facilitates large-scale citizen-science research. This approach involves the recruitment of volunteer participants, who typically complete experiments because the experiments are intrinsically rewarding, with larger and more diverse samples than are typically feasible with in-laboratory research^86, 108^. A total of 68,206 participants began the experiment, the first in January 2019 and the last in October 2021. Demographics in the sub-sample of United States participants are in Extended Data Table 7.

We played participants vocalizations from a subset of the corpus, excluding those that were less than 10 seconds in duration (*n* = 111) and those with confounding sounds produced by a source other than the target voice in the first 5 seconds of the recording (e.g., a crying baby or laughing adult in the background; *n* = 366), as determined by two independent annotators who remained unaware of vocalization type and fieldsite with disagreements resolved by discussion. A test of the robustness of the main effects to this exclusion decision is in SI Text 1.4. We also excluded participants who reported having previously participated in the same experiment (*n* = 3,889); participants who reported being younger than 12 years old (*n* = 1,519); and those who reported having a hearing impairment (*n* = 1,437).

This yielded a sample of 51,065 participants (gender: 22,862 female, 27,045 male, 1,117 other, 41 did not disclose; age: median 22 years, interquartile range 18-29). Participants self-reported living in 187 different countries (Fig. 3b) and self-reported speaking 172 first languages and 147 second languages (27 of which were not in the list of first languages), for a total of 199 different languages. Roughly half the participants were native English speakers from the United States. We supplemented these data with a paid online experiment, to increase the sampling of a subset of recordings in the corpus (SI Text 1.8).

Participants listened to at least 1 and at most 16 vocalizations drawn from the subset of the corpus (as they were free to leave the experiment before completing it) for a total of 495,512 ratings (infant-directed song: *n* = 139,708; infant-directed speech: *n* = 99,482; adult-directed song: *n* = 132,124; adult-directed speech: *n* = 124,198). The vocalizations were selected with blocked randomization, such that a set of 16 trials included 4 vocalizations in English and 12 in other languages; this method ensured that participants heard a substantial number of non-English vocalizations. This yielded a median of 516.5 ratings per vocalization (interquartile range 315-566; range 46-704) and thousands of ratings for each society (median = 22,974; interquartile range 17,458-25,177). The experiment was conducted only in English, so participants likely had at least a cursory knowledge of English; a test of the robustness of the main effects when excluding English-language recordings is in SI Text 1.4.

We asked participants to classify each vocalization as either directed toward a baby or an adult. The prompt “Someone is speaking or singing. Who do you think they are singing or speaking to?” was displayed while the audio played; participants could respond with either “adult” or “baby”, either by pressing a key corresponding to a drawing of an infant or adult face (when the participant used a desktop computer) or by tapping one of the faces (when the participant used a tablet or smartphone). The locations of the faces (left vs. right on a desktop; top vs. bottom on a tablet or smartphone) were randomized participant-wise. Screenshots are in Extended Data Fig. 4.

We asked participants to respond as quickly as possible, a common instruction in perception experiments, to reduce variability that could be introduced by participants hearing differing lengths of each stimulus; to reduce the likelihood that participants used linguistic content to inform their decisions; and to facilitate a response-time analysis (Extended Data Fig. 8), as jsPsych provides reliable response time data^109^. We also used the response time data as a coarse measure of compliance, by dropping trials where participants were likely inattentive, responding very quickly (less than 500 ms) or slowly (more than 5 s). Most response times fell within this time window (82.1% of trials).

The experiment included two training trials, using English-language recordings of a typically infant-directed song (“The wheels on the bus”) and a typically adult-directed song (“Hallelujah”); 92.7% of participants responded correctly by the first try and 99.5% responded correctly by the second try, implying that the vast majority of the participants understood the task.

As soon as they made a choice, playback stopped. After each trial, we told participants whether or not they had answered correctly and how long, in seconds, they took to respond. At the end of the experiment, we showed participants their total score and percentile rank (relative to other participants).

## End notes

### Data, code, and materials availability

A reproducible R Markdown manuscript; data, analysis code, and visualizations; supplementary fieldsite- level data; the recording collection protocol; and code for the naïve listener experiment are available at https://github.com/themusiclab/infant-speech-song. The audio corpus is available at https://doi.org/10.5281/zenodo.5525161. The preregistration for the auditory analyses is at https://osf.io/5r72u. Readers may participate in the naïve listener experiment by visiting https://themusiclab.org/quizzes/ids.

## Acknowledgments

This research was supported by the Harvard University Department of Psychology (M.M.K. and S.A.M.); the Harvard College Research Program (H.L-R.); the Harvard Data Science Initiative (S.A.M.); the National Institutes of Health Director’s Early Independence Award DP5OD024566 (S.A.M. and C.B.H.); the Academy of Finland Grant 298513 (J. Antfolk); the Royal Society of New Zealand Te Apārangi Rutherford Discovery Fellowship RDF-UOA1101 (Q.D.A., T.A.V.); the Social Sciences and Humanities Research Council of Canada (L.K.C.); the Polish Ministry of Science and Higher Education grant N43/DBS/000068 (G.J.); the Fogarty International Center (P.M., A. Siddaiah, C.D.P.); the National Heart, Lung, and Blood Institute, and the National Institute of Neurological Disorders and Stroke Award D43 TW010540 (P.M., A. Siddaiah); the National Institute of Allergy and Infectious Diseases Award R15-AI128714-01 (P.M.); the Max Planck Institute for Evolutionary Anthropology (C.T.R., C.M.); a British Academy Research Fellowship and Grant SRG-171409 (G.D.S.); the Institute for Advanced Study in Toulouse, under an Agence nationale de la recherche grant, Investissements d’Avenir ANR-17-EURE-0010 (L.G., J. Stieglitz); the Fondation Pierre Mercier pour la Science (C.S.); and the Natural Sciences and Engineering Research Council of Canada (S.E.T.). We thank the participants and their families for providing recordings; Lawrence Sugiyama for supporting pilot data collection; Juan Du, Elizabeth Pillsworth, Polly Wiessner, and John Ziker for collecting or attempting to collect additional recordings; Ngambe Nicolas for research assistance in the Republic of the Congo; Zuzanna Jurewicz for research assistance in Toronto; Maskota Delfi and Rustam Sakaliou for research assistance in Indonesia; Willy Naiou and Amzing Altrin for research assistance in Vanuatu; S. Atwood, Anna Bergson, Dara Li, Luz Lopez, and Emilė Radytė for project-wide research assistance; and Jonathan Kominsky, Lindsey Powell, and Lidya Yurdum for feedback on the manuscript.

## Author contributions

- S.A.M. and M.M.K. conceived of the research, provided funding, and coordinated the recruitment of collaborators and creation of the corpus.
- S.A.M. and M.M.K. designed the protocol for collecting vocalization recordings with input from D.A., who piloted it in the field.
- L.G., A.G., G.J., C.T.R., M.B.N., A. Martin, L.K.C., S.E.T., J. Song, M.K., A. Siddaiah, T.A.V., Q.D.A., J. Antfolk, P.M., A. Schachner, C.D.P., G.D.S., S.K., M.S., S.A.C., J.Q.P., C.S., J. Stieglitz, C.M., R.R.S., and B.M.W collected the field recordings, with support from E.A., A. Salenius, J. Andelin, S.C.C., M.A., and A. Mabulla.
- S.A.M., C.M.B., and J. Simson designed and implemented the online experiment.
- C.J.M. and H.L-R. processed all recordings and designed the acoustic feature extraction with S.A.M. and M.M.K.; C.M.B. provided associated research assistance.
- C.M. designed the fieldsite questionnaire with assistance from M.B. and C.J.M., who collected the data from the principal investigators.
- C.B.H. and S.A.M. led analyses, with additional contributions from C.J.M., M.B., D.K., and M.M.K.
- C.B.H. and S.A.M. designed the figures.
- C.B.H. wrote computer code, with contributions from S.A.M., C.J.M., and M.B.
- D.K. conducted code review.
- C.J.M., H.L-R., M.M.K., and S.A.M. wrote the initial manuscript.
- C.B.H. and S.A.M. wrote the first revision, with contributions from C.J.M. and M.B.
- S.A.M. wrote the second and third revisions, with contributions from C.B.H. and C.J.M.

## Ethics

Ethics approval for the collection of recordings was provided by local institutions and/or the home institution of the collaborating author who collected data at each fieldsite. These included the Bioethics Committee, Jagiellonian University (1072.6120.48.2017); Board for Research Ethics, Åbo Akademi University; Committee on the Use of Human Subjects, Harvard University (IRB16-1080 and IRB18-1739); Ethics Committee, School of Psychology, Victoria University of Wellington (0000023076); Human Investigation Committee, Yale University (MODCR00000571); Human Participants Ethics Committee, University of Auckland (018981); Human Research Protections Program, University of California San Diego (161173); Institutional Review Board, Arizona State University (STUDY00008158); Institutional Review Board, Florida International University (IRB-17-0067); Institutional Review Board, Future Generations University; Max Planck Institute for Evolutionary Anthropology; Research Ethics Board, University of Toronto (33547); Research Ethics Committee, University College London (13121/001); Review Board for Ethical Standards in Research, Toulouse School of Economics/IAST (2017-06-001 and 2018-09-001); and Tanzania Commission for Science and Technology (COSTECH). Ethics approval for the naïve listener experiment was provided by the Committee on the Use of Human Subjects, Harvard University (IRB17-1206). Informed consent was obtained from all participants.

## Additional information

The authors declare no competing interests.

**Supplementary information** is available for this paper.

**Correspondence and requests for materials** should be addressed to C.B.H., C.J.M., and S.A.M.

## Supplementary Text

### 1.1 Technical details of the acoustic feature extraction

We extracted acoustic features with four sets of tools, described below, and also preprocessed them to reduce the influence of atypical observations.

#### 1.1.1 Praat

We extracted intensity, pitch, and first and second formant values from the denoised recordings every 0.03125 seconds. For female participants, the pitch floor was set at 100 Hz, with a pitch ceiling at 600 Hz, and a maximum formant of 5500 Hz. For male participants, these values were 75 Hz, 300 Hz, and 5000 Hz, respectively. From these data, several summary values were calculated for each recording: mean and maximum first and second formants, mean pitch, and minimum intensity. In addition to these summary statistics, we measured the intensity and pitch rates as change in these values over time. For vowel measures, the first and second formants were used to calculate both the average vowel space used, as well as the vowel change rate (measured as change in Euclidean formant space) over time.

#### 1.1.2 MIRtoolbox

All MIRtoolbox (v. 1.7.2) features were extracted with default parameters^102^. *mirattackslope* returns a list of all attack slopes detected, so final analyses were done on summary features (e.g., mean, median, etc.). Final analyses were also done on summary features for *mirroughness*, which returns time series data of roughness measures in 50ms windows. We RMS-normalized the mean of *mirroughness*, following previous work^110^. MIRtoolbox features were computed on the denoised recordings, with the exception of *mirtempo* and *mirpulseclarity*, where removing the silences between vocalizations would have altered the tempo.

#### 1.1.3 Rhythmic variability

For temporal modulation spectra we followed a previous method^111^, which combines discrete Fourier transforms applied to contiguous six-second excerpts. To analyze the entirety of each recording, we appended all recordings with silence to be exact multiples of six-seconds. The location of the peak (Hz) and variance of the temporal modulation spectra were extracted from their RMS values. Because intervening silence would influence temporal modulation measures, we computed them on recordings *before* they had been denoised.

#### 1.1.4 Normalized pairwise variability index (nPVI)

The nPVI represents the temporal variance of data with discrete events, which makes it especially useful for comparing speech and music^103^. We used an automated syllable- and phrase-detection algorithm to extract events^104^. We computed nPVI in two ways: by averaging the nPVI of each phrase within a recording, as well as by treating the entire recording as a single phrase. Because intervening silence would influence nPVI measures, we computed them on recordings *before* they had been denoised.

#### 1.1.5 Preprocessing

Automated acoustic analyses are highly sensitive at extremes (e.g., impossible values caused by non-vocal sounds, like loud wind). To correct for these issues, we Winsorized all acoustic variables. This process defines observations exceeding the lowest and highest 5 percentile ranks as outliers, recoding them as the values of those percentile boundaries. These data were used for all acoustic analyses. This approach is generally preferable to trimming extreme values, as trimming overcompensates for outliers by removing them entirely^112^.

Analyses of the acoustic features using an alternate method (i.e., imputing extreme values with the mean observation for each feature within each fieldsite) yielded comparable results; readers are welcome to try alternate trimming methods with the open data and materials.

In the cases of three acoustic features (roughness, vowel travel rate, and pulse clarity), we used log-transformed data, because the raw data were highly skewed. This decision was supported by the exploratory-confirmatory approach; that is, results replicated across both exploratory and confirmatory samples in the log-transformed data.

### 1.2 Alternate analysis of acoustic features via principal-components approach

We conducted an exploratory principal components analysis of the full 94 acoustic variables (Extended Data Fig. 3). The analysis accounted for ∼40% of total variability in acoustic features. The results provide convergent evidence that the main forms of acoustic variation partition into orthogonal clusters that most strongly distinguish speech from song overall (in PC1); most strongly distinguish infant-directedness in *song* (in PC2); and most strongly distinguish infant-directedness in *speech* (in PC3). Factor loadings are in Extended Data Table 8; these largely corroborate the findings of the LASSO and exploratory-confirmatory analyses.

One further pattern that the principal components analysis highlights is that infant-directedness makes speech more “songlike”, in terms of higher pitch and reduced roughness (PC3); but speech strongly differed from song overall in terms of the variability and rate of variability of pitch, intensity, and vowels, and infant-directedness further exaggerated these differences for speech (PC1).

### 1.3 Quantifying sensitivity with signal detection theory

To quantify the listener sensitivity to infant-directedness in speech and song, and to quantify their response biases, we computed the metrics of *d^′^* and *c* (*criterion*) over the stimuli. These quantities were calculated with standard techniques from signal detection theory^113^.

Specifically, a response on a given trial was coded as a hit if the trial was an infant-directed vocalization and the participant correctly responded with baby; a miss if for an infant-directed vocalization, they responded adult; a false-alarm if for an adult-directed vocalization, they responded baby; and a correct-reject if for an adult-directed vocalization, they correctly responded adult.

The hit rate *H* was then computed as the total number of hits for a given recording, divided by the total number of hits plus the misses; the false-alarm rate *F* was computed as the total number of misses for a given recording, divided by the total number of false-alarms plus the correct-rejects. These scores were then conservatively adjusted with the log-linear correction for extreme scores^114^, and finally *d^′^* was estimated via the following equation, where the function *z*(*·*) represents the inverse of the normal cumulative distribution function:

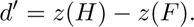

Criterion (*c*) was estimated as:

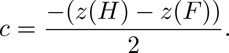

### 1.4 Robustness tests of main results in naïve listener experiment

On the suggestion of an anonymous reviewer, we repeated the main analyses of the naïve listener experiment (i.e., estimated sensitivity to infant-directedness in speech and song) with two alternate data exclusion strategies. First, the analyses and figures in the main text only study ratings of recordings that contained minimal extraneous sounds (such as a baby crying; see Methods). To ensure that the exclusion of these recordings did not account for the main findings, we repeated the analyses while including ratings of *all* recordings, including those with putatively confounding background sounds. They robustly replicated, with comparable effect sizes (speech: *d^′^* = 1.13, 95% CI [0.48, 1.77]; song: *d^′^* = 0.54, 95% CI [0.23, 0.86]; *ps < .*05).

A further potential confound concerns listeners’ familiarity with the languages spoken or sung in the recordings. In the main text analyses, we explicitly model the expected differences in sensitivity that could result from lower or higher degrees of linguistic relatedness between the vocalizer and the listener (see, e.g., Fig. 3c). However, because the experiment was only conducted in English, many participants likely could understand at least some parts of the English-language vocalizations. To ensure that these recordings did not account for the main findings, we repeated the analyses while excluding all English-language recordings. These recordings came predominantly from the Wellington, San Diego, and Toronto fieldsites (where nearly all recordings were in English) but also appeared elsewhere, such as the Arawak fieldsite (where English Creole recordings were often comprehensible to English speakers), and in a few other sites, when a speaker happened to be bilingual and produce English-language vocalizations. The results robustly replicated with these exclusions (speech: *d^′^*= 0.83, 95% CI [0.33, 1.33]; song: *d^′^* = 0.33, 95% CI [0.08, 0.57]; *ps < .*05).

### 1.5 Demographic analyses of a subsample of naïve listeners

An anonymous reviewer raised the possibility that conducting the naïve listener experiment online, as opposed to in a laboratory, reduced the diversity of the sample; if so, this could bias the results of the experiment, in principle. To test this question, we analyzed demographic information from participants living in the United States, who provided income, education level, and ethnicity data.

Descriptive statistics revealed that the subsample of United States participants was highly diverse (Extended Data Table 7), including, for example, representation from all ethnicity categories currently defined by the National Institutes of Health, and a broad range of annual household incomes. The sample was generally more representative of the United States population than are samples recruited in typical laboratory studies, which may skew towards wealthier samples with representation of fewer ethnicity categories^107, 108^.

Nevertheless, we proceeded by asking whether demographic factors were likely to affect people’s ability to perceive infant-directedness. We ran mixed-effect regressions for each of the available demographic variables with random intercepts for the vocalist in the recording, and fixed effects for vocalization type and the demographic factor. While the main effects of income, education, or race on task performance were statistically significant (*ps* < 0.0001), in all cases, the effect sizes were tiny, explaining ∼0.1% of variance in the model. These findings imply that the choice of a citizen-science approach likely did not bias the results of the experiment, at least in United States participants.

### 1.6 Society-level predictors for naïve listener data

Listener sensitivity within each fieldsite was correlated with a number of society-level characteristics: rank-order population size (speech: *τ* = 0.51; song: *τ* = 0.58), distance from fieldsite to nearest urban center (speech: *r* = -0.78; song: *r* = -0.51), and number of children per family (speech: *r* = -0.53; song: *r* = -0.72; all *ps < .*001). Each of these predictors were highly correlated with each other (all *r* > 0.6), however, suggesting that they did not each contribute unique variance. There was no correlation with ratings of how frequently infant-directed vocalizations were used within each society (*ps > .*4). These findings suggest that at least some cross-fieldsite variability in listener sensitivity to infant-directedness is attributable to the *cultural* relatedness between vocalizers and listeners (as opposed to the *linguistic* relatedness analyzed in in the Main Text and Fig. 3c).

### 1.7 Simulated infant-directed vocalizations

Prior research has shown that simulated infant-directedness is qualitatively similar, albeit less exaggerated than when authentic, for both speech^115^ and song^35^. Indeed, a model of the naïve listener results adjusting for fieldsite indeed showed a small decrease in “baby” guesses when an infant was not present (ID song: 6.4%, ID speech: 7.5%, AD song: -6.5%, AD speech: -4.2%, *p*s < .0001), but this effect was not stronger for vocalizations that were infant-directed compared to adult-directed (*χ*^2^(1) = 2.93, *p* = 0.09). Both the naive listener results and acoustic analyses were robust to whether these simulated infant-directed vocalizations were included or excluded, however, implying that the use of simulated infant-directed vocalizations did not undermine the robustness of the main effects.

### 1.8 Additional data collection via Prolific

In revising this manuscript, we discovered that a small subset of the corpus had been erroneously excluded from the main experiment. In most cases, these were recordings that had been too-conservatively edited to be too short to include in the experiment (but could reasonably be edited to include longer sections of audio); in some other cases, the original excerpting included confounding background noises that, upon additional editing, were avoidable. To ensure maximal coverage of the fieldsites studied here, we re-excerpted the audio of 103 examples and collected supplemental naïve listener data on these recordings via a Prolific experiment (N = 97, 54 male, 42 female, 1 other, mean age = 29.7 years). The Prolific experiment was identical to the citizen-science experiment, except that each participant was paid US$15/hr, rather than volunteering; and each participant rated 188 recordings instead of up to 16.

We included in the Prolific experiment the set of recordings that were erroneously excluded from the citizen-science experiment, along with 85 additional recordings randomly selected from those that *were* included in the citizen-science experiment, so as to ensure that each Prolific participant heard a balanced set of vocalization types. The two cohorts’ ratings of the recordings in common across the two experiments were highly correlated (*r* = 0.95, *p* < 0.0001), demonstrating that they had similar intuitions concerning infant-directedness in speech and song. As such, in the main text, we report all the ratings together without disambiguating between the cohorts.

---

**Extended Data Fig. 1.**
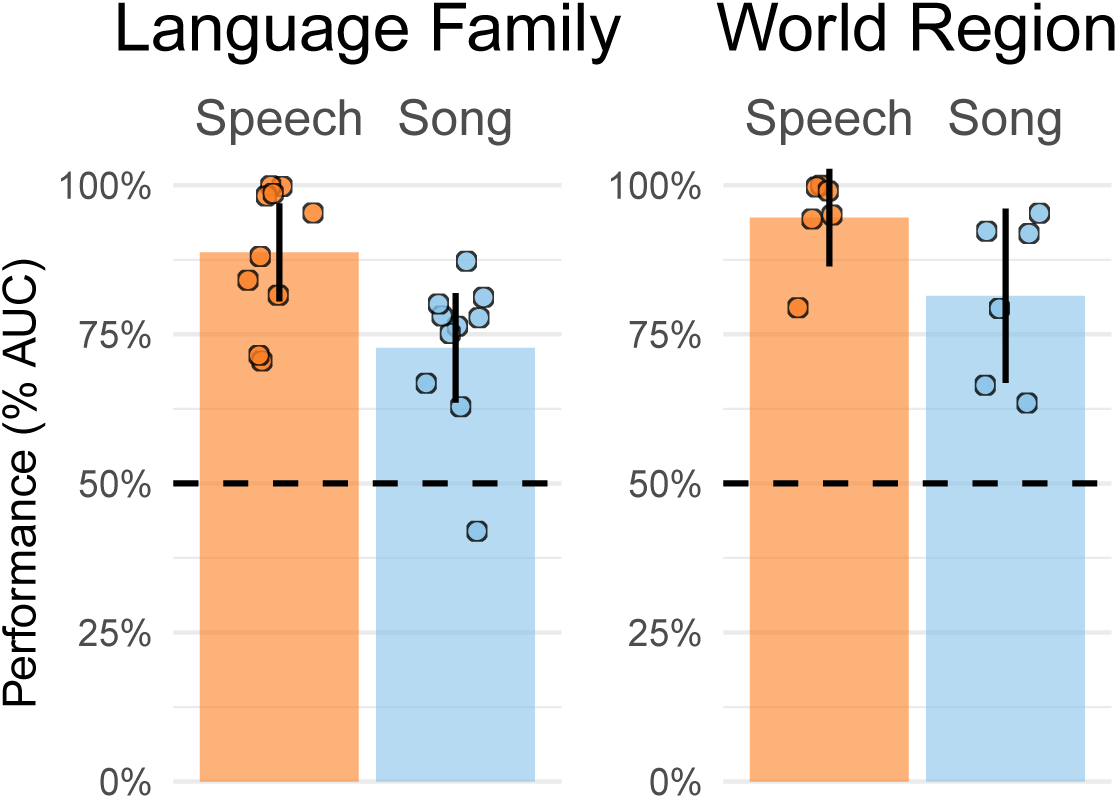
LASSO classification of acoustic features with alternate cross-validation approaches. We repeated the main LASSO analysis (Fig. 1b) twice, but rather than conducting *k*-fold cross-validation across fieldsites, we did so across language families and world regions (see descriptive information about the fieldsites in Table 1). The results replicated robustly across both models, with corpus-wide classification performance significantly above chance in all cases. The vertical bars represent the overall classification performance (quantified via receiver operating characteristic/area under the curve; AUC); the error bars represent 95% confidence intervals; the points represent the performance estimate for each language family or world region; and the horizontal dashed lines indicate chance level of 50% AUC.

**Extended Data Fig. 2.**
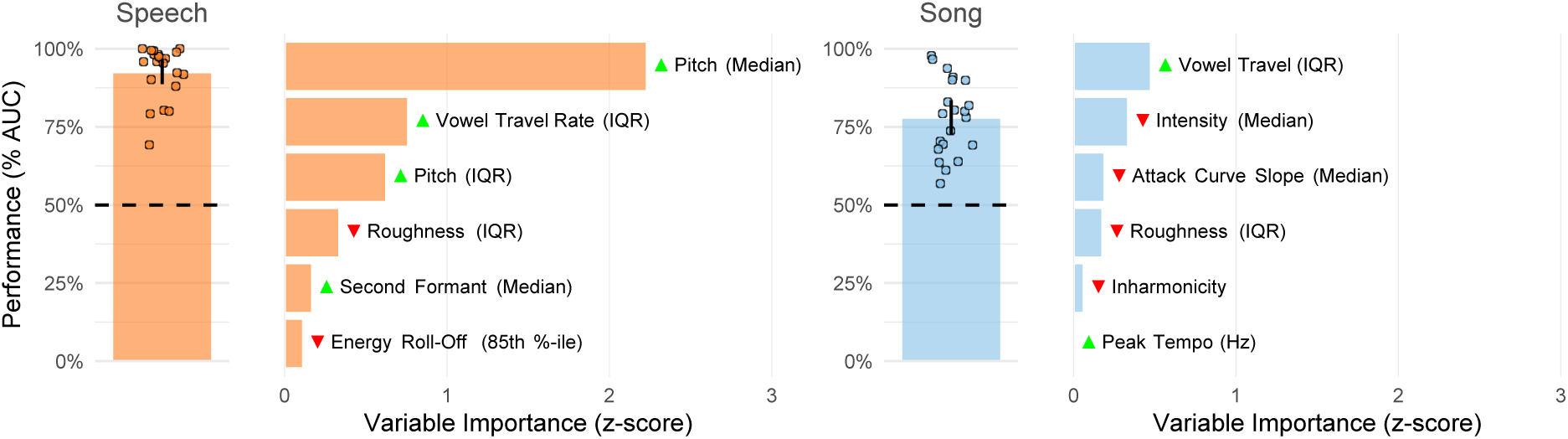
Replication of main LASSO results using unedited audio. As a test of robustness, we repeated the main LASSO analyses (Fig. 1b) with acoustic features extracted from raw, unedited audio. This approach ensures that the main results are not attributable to idiosyncrasies in the audio introduced by the editing process. The results repeated robustly, with above-chance performance in all fieldsites for both speech and song, and with the 3 most influential acoustic features selected by the model repeating across both specifications (see Fig. 1b). The vertical bars represent the overall classification performance (quantified via receiver operating characteristic/area under the curve; AUC); the error bars represent 95% confidence intervals; the points represent the average performance for each fieldsite; and the horizontal dashed lines indicate chance level of 50% AUC. The horizontal bars show the acoustic characteristics with the largest influence in each classifier; the green and red triangles indicate the direction of the effect, e.g., with median pitch having a large, positive effect on classification of infant-directed speech. See Methods for further details.

**Extended Data Fig. 3.**
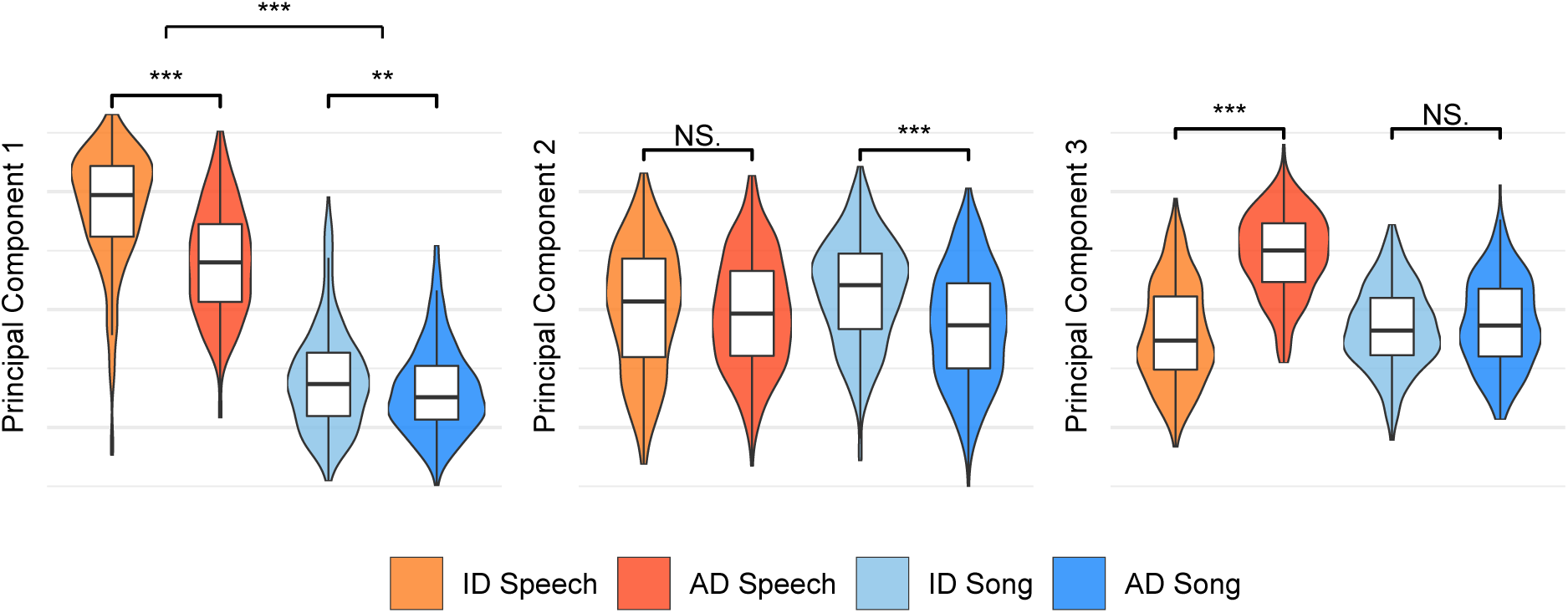
Principal-components analysis of acoustic features. As an alternative approach to the acoustics data, we ran a principal-components analysis on the full 94 acoustic variables, to test whether an unsupervised method also yielded opposing trends in acoustic features across the different vocalization types. It did. Three components explained approximately 40% of total variability in the acoustic features. Moreover, the clearest differences between vocalization types accorded with the LASSO and mixed-effects modeling (Figs. 1b and 2). The first principal component most strongly differentiated speech and song, overall; the second most strongly differentiated infant-directed song from adult-directed song; and the third most strongly differentiated infant-directed speech from adult-directed speech. The violins indicate kernel density estimations and the boxplots represent the medians and interquartile ranges. Significance values are computed via Wilcoxon signed-rank tests; \**p <* 0.05, \*\**p <* 0.01, \*\*\**p <* 0.001. Feature loadings are in Extended Data Table 8.

**Extended Data Fig. 4.**
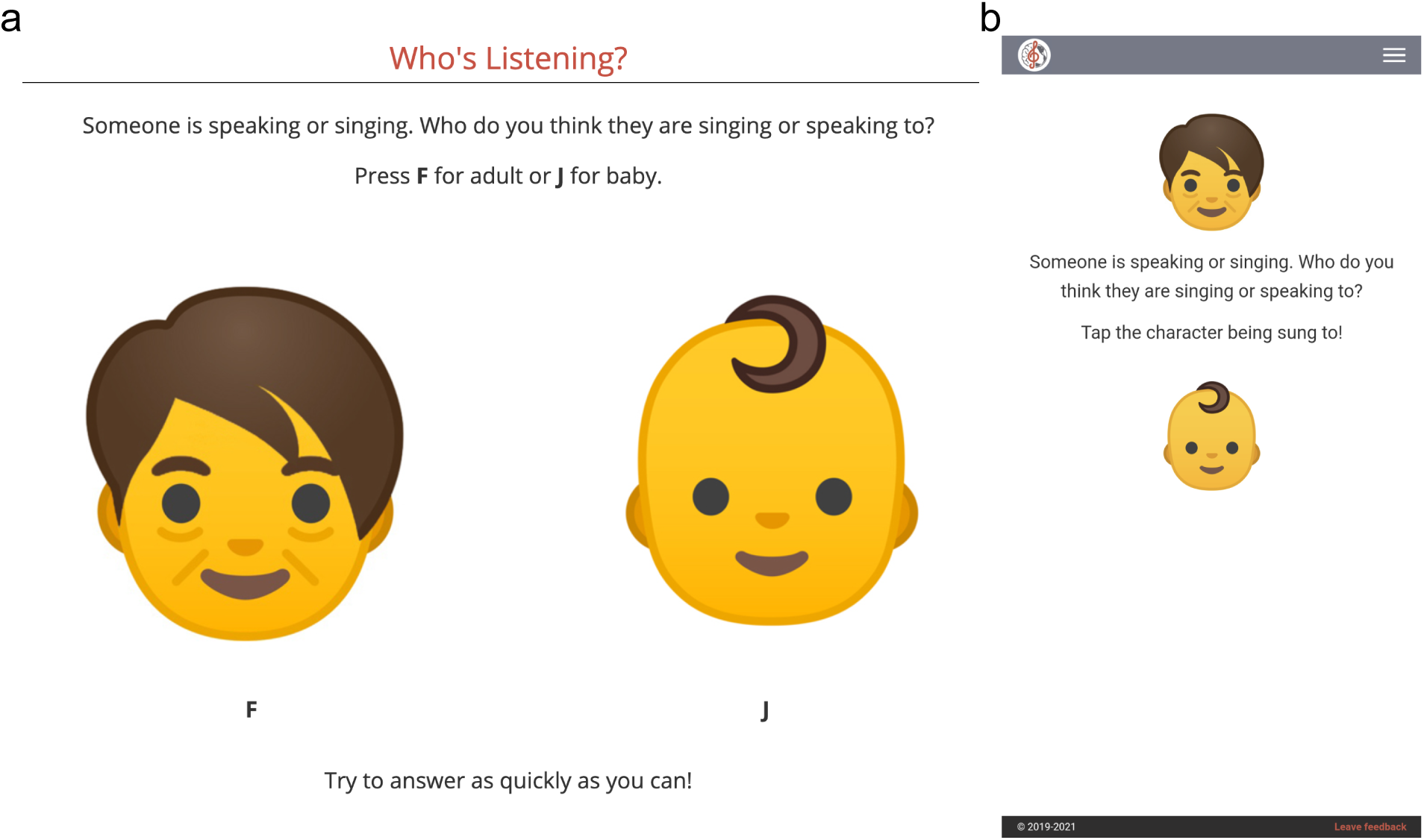
Screenshots from the naïve listener experiment. On each trial, participants heard a randomly selected vocalization from the corpus and were asked to quickly guess to whom the vocalization was directed: an adult or a baby. The experiment used large emoji and was designed to display comparably on desktop computers (**a**) or tablets/smartphones (**b**).

**Extended Data Fig. 5.**
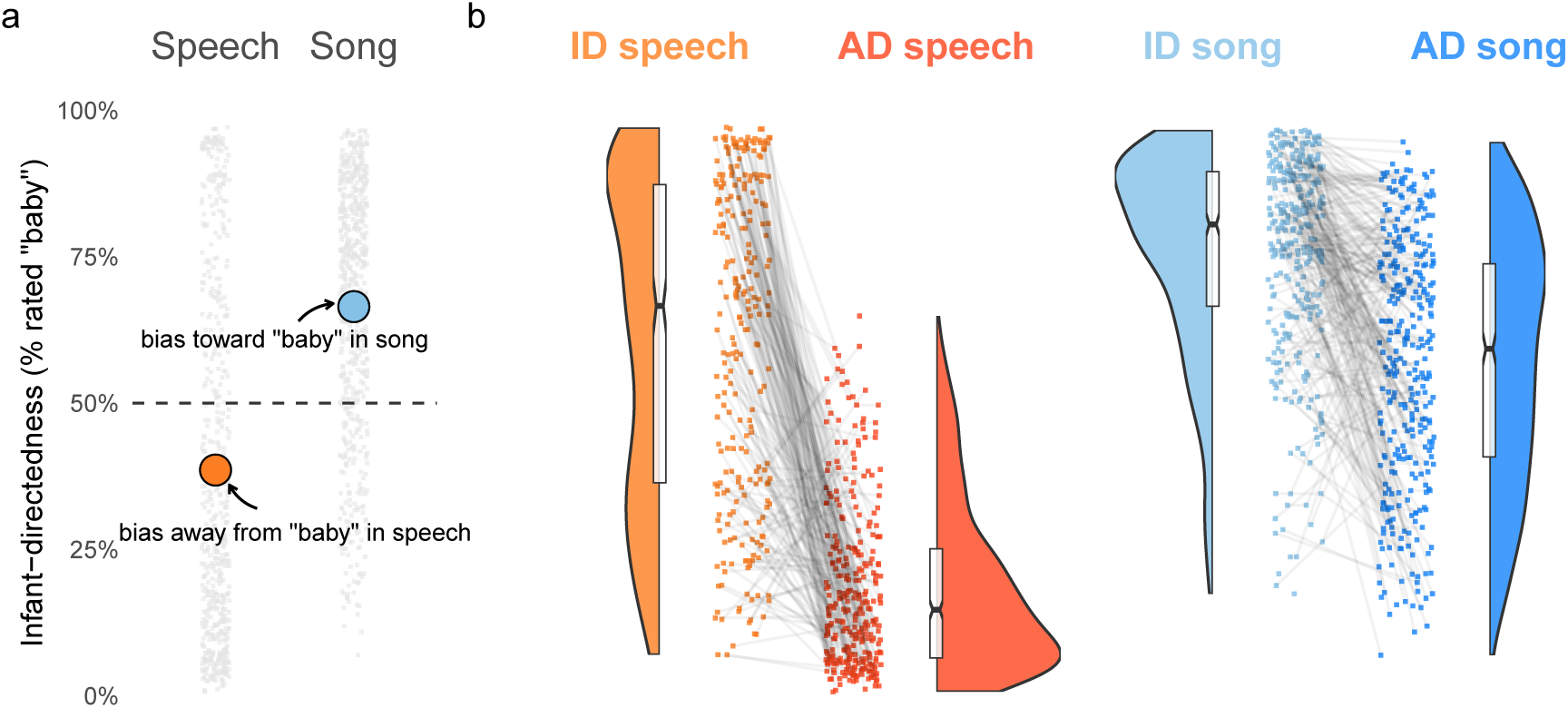
Response biases in the naïve listener experiment. **a**, Listeners showed reliable biases: regardless of whether a vocalization was infant- or adult-directed, the listeners gave speech recordings substantially fewer “baby” responses than expected by chance, and gave song recordings substantially more “baby” responses. The gray points represent average ratings for each of the 1615 recordings in the corpus, split by speech and song; the orange and blue points indicate the means of each vocalization type; and the horizontal dashed line represents hypothetical chance level of 50%. **b**, Despite the response biases, within speech and song, the raw data nevertheless showed clear differences between infant-directed and adult-directed vocalizations, i.e., by comparing infant-directedness scores within the same voice, across infant-directed and adult-directed vocalizations (visible here in the steep negative slopes of the gray lines). The main text results report only *d^′^* statistics for these data, for simplicity, but the main effects are nonetheless visible here in the raw data. The points indicate average ratings for each recording; the gray lines connecting the points indicate the pairs of vocalizations produced by the same voice; the half-violins are kernel density estimations; the boxplots represent the medians, interquartile ranges, and 95% confidence intervals (indicated by the notches); and the horizontal dashed lines indicate the response bias levels (from **a**).

**Extended Data Fig. 6.**
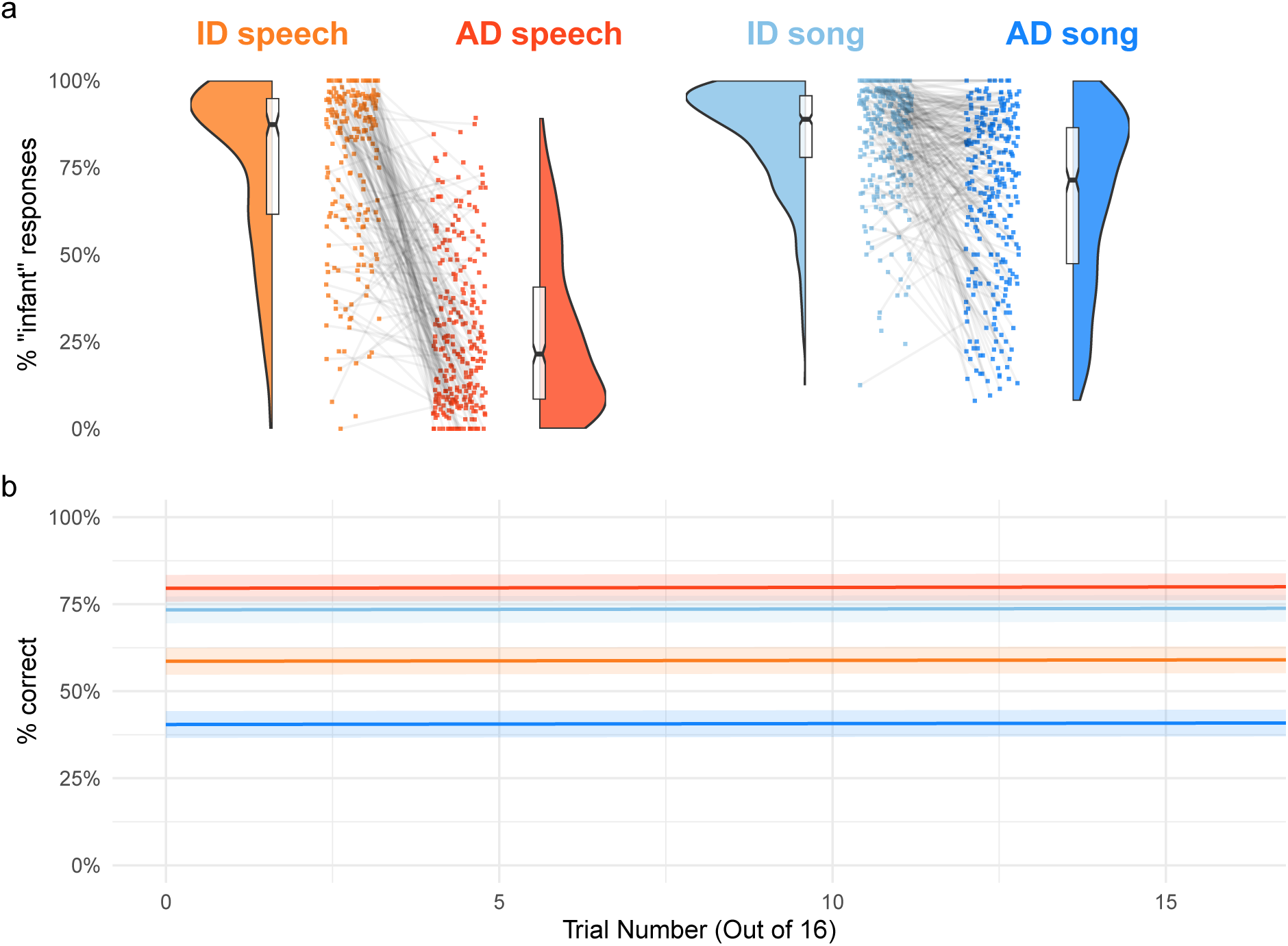
The main effects in the naïve listener experiment are not attributable to learning. **a**, This panel repeats the raw accuracy data reported in Extended Data Fig. 5b, but using only data from responses that were participants’ first trial, to avoid the possibility of any learning effects over the course of their participation. The results do not change appreciably. See further details in the caption to Extended Data Fig. 5. **b**, Over the course of multiple trials in the experiment, which contained corrective feedback, participants’ raw accuracy barely increased. The lines depict linear regressions for each of the four vocalization types and the shaded regions depict 95% confidence intervals.

**Extended Data Fig. 7.**
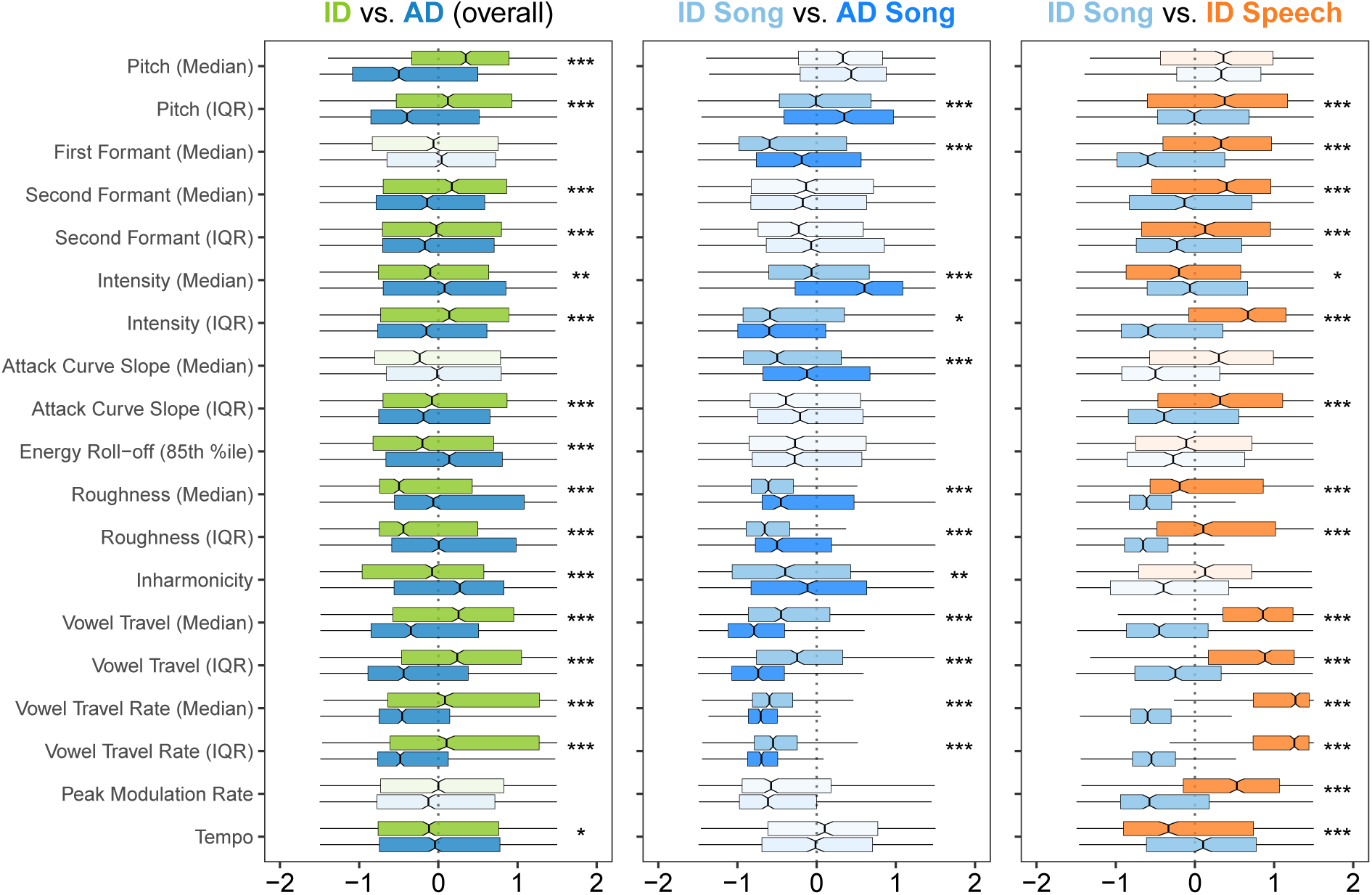
Exploratory-confirmatory selected acoustic features for pre-registered analyses. The preregistered analyses included comparisons of the acoustic features of infant-directed vocalizations, regardless of whether they included speech or song. For the reasons discussed in the Methods, and per the results reported in Fig. 2, these results should be interpreted with caution, as direct comparisons of acoustic features across modalities (language vs. music) may be spurious or may hide underlying variation within each modality. Moreover, these analyses do not include fieldsite-level random effects, so they are less conservative than those reported in Fig. 2 (i.e., they identify a larger number of acoustic features). The boxplots show the 25 acoustic features with a significant difference in at least one main comparison (e.g., infant-directed song vs. infant-directed speech, in the right panel), in both the exploratory and confirmatory analyses. All variables are normalized across participants. The boxplots represent the median and interquartile range; the whiskers indicate 1.5 *×* IQR; and the notches represent the 95% confidence intervals of the medians. Faded comparisons did not reach significance in exploratory analyses. Significance values are computed via linear combinations, following mixed-effects models; \**p <* 0.05, \*\**p <* 0.01, \*\*\**p <* 0.001. Prespecified hypotheses about each comparison are posted in the project GitHub repository.

**Extended Data Fig. 8.**
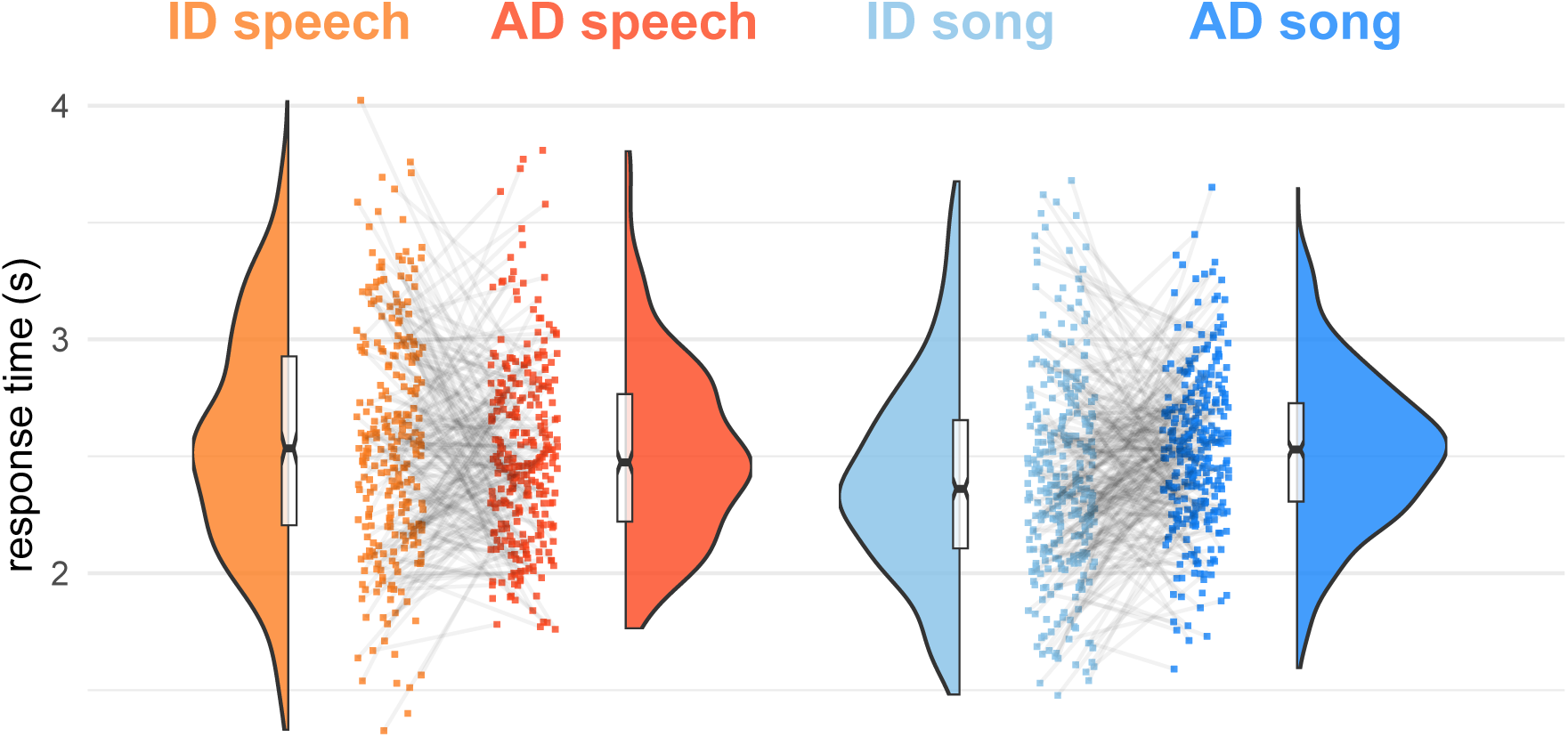
Response-time analysis of naïve listener experiment. We recorded the response times of participants in their mobile or desktop browsers, using jsPsych (see Methods), and asked whether, when responding correctly, participants more rapidly detected infant-directedness in speech or song. They did not: a mixed-effects regression predicting the difference in response time between infant-directed and adult-directed vocalizations (within speech or song), adjusting hierarchically for fieldsite and world-region, yielded no significant differences (*p*s > .05 from linear combination tests). The points indicate median response times for each recording across all correct responses; the gray lines connecting the points indicate the pairs of vocalizations produced by the same participant; the half-violins are kernel density estimations; and the boxplots represent the medians, interquartile ranges, and 95% confidence intervals (indicated by the notches).

**Extended Data Table 1.**
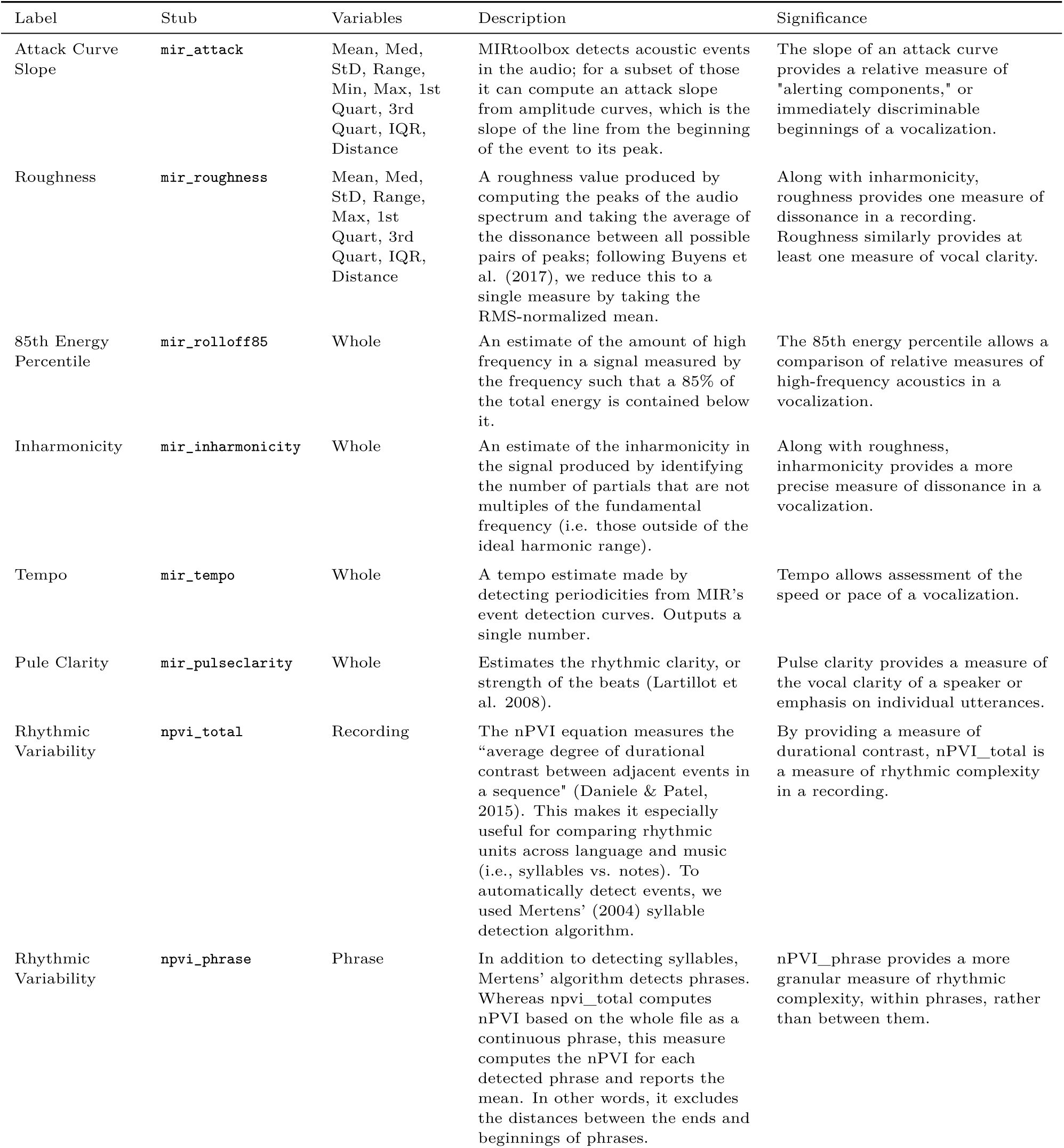
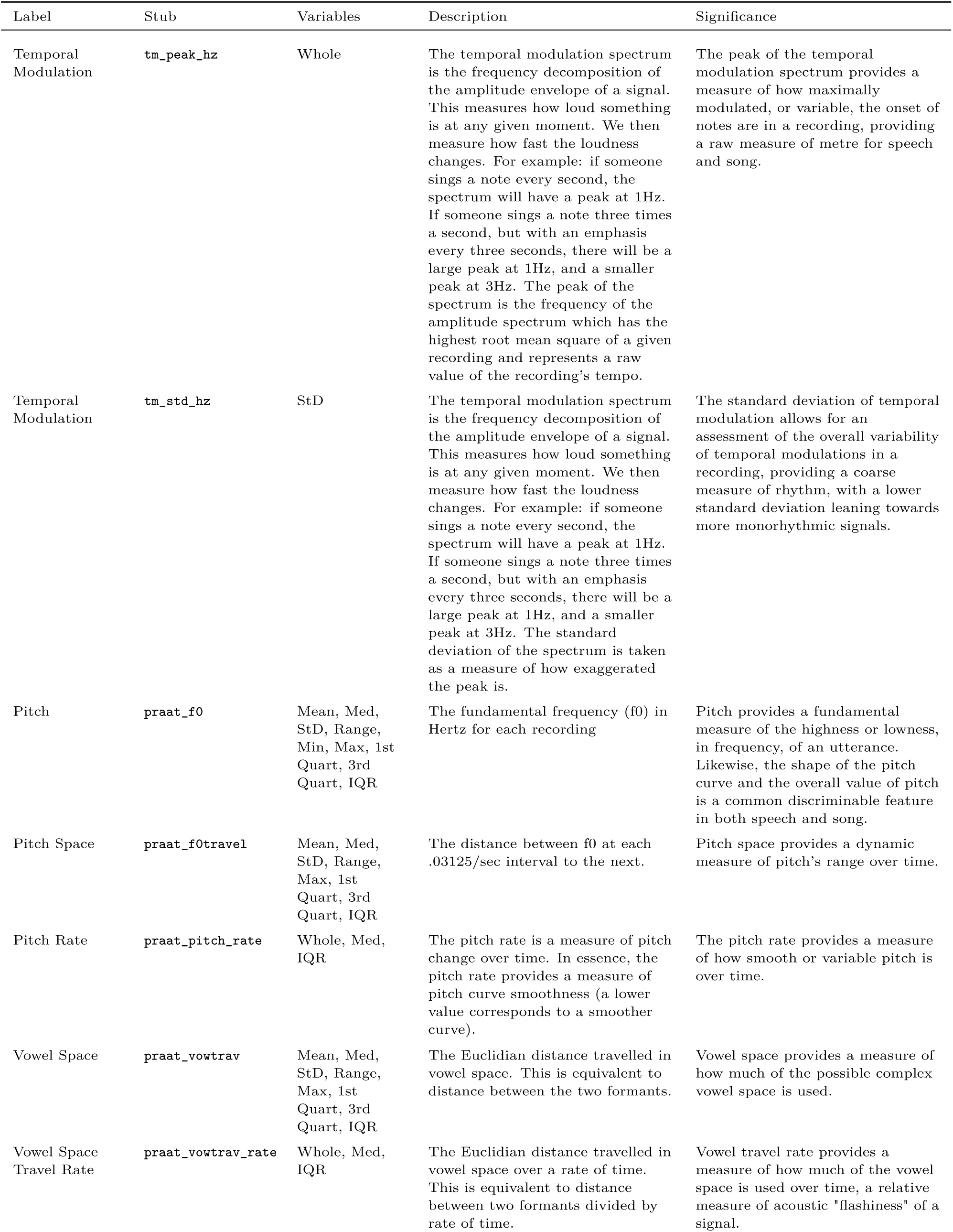
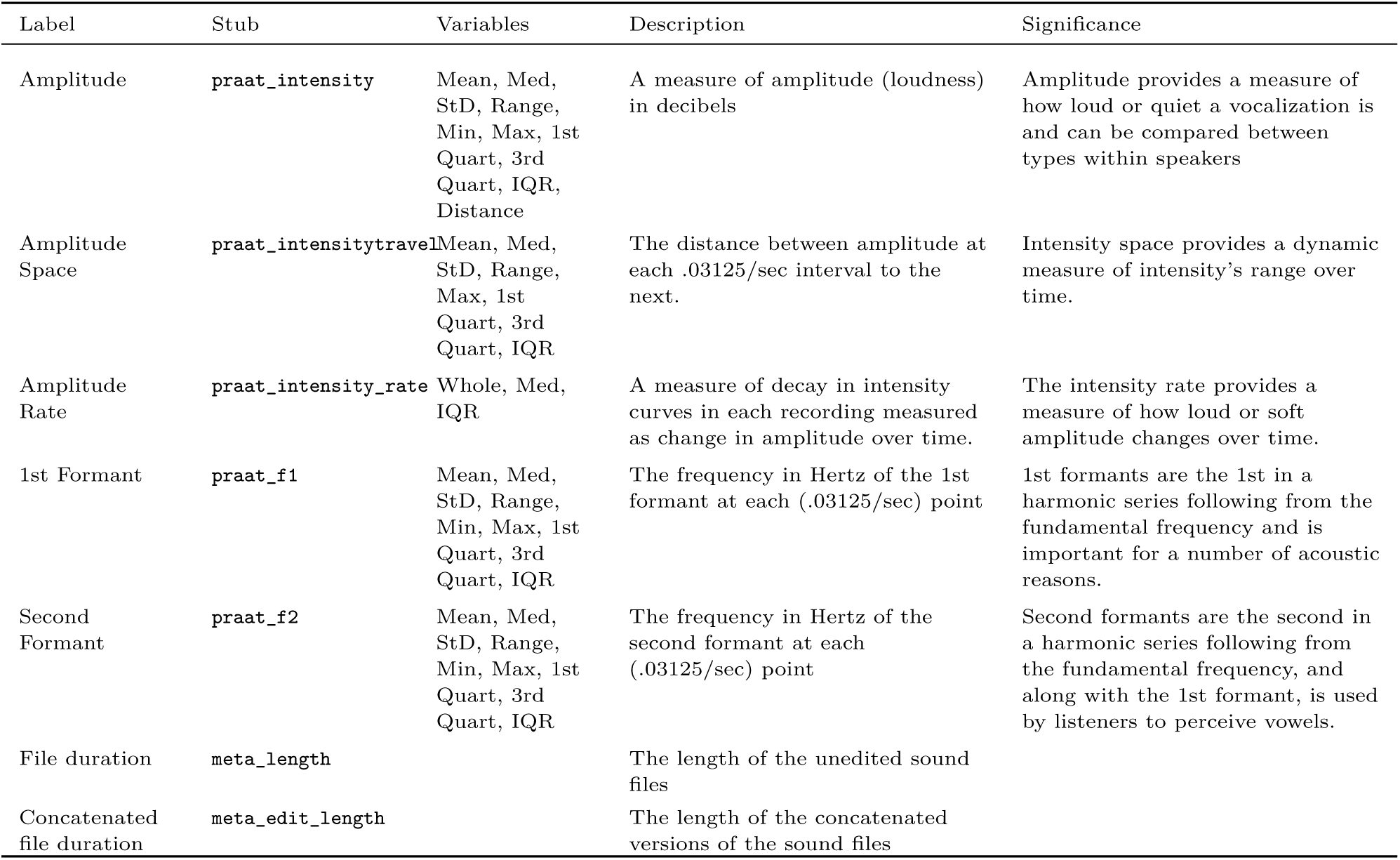
Codebook for acoustic features. Variable names are stubs; in the datasets on the project GitHub repository, suffixes are added to denote summary statistics (e.g., mir_attack_mean).

| Label | Stub | Variables | Description | Significance |
| --- | --- | --- | --- | --- |
| Attack Curve Slope | `mir_attack` | Mean, Med, StD, Range, Min, Max, 1st Quart, 3rd Quart, IQR, Distance | MIRtoolbox detects acoustic events in the audio; for a subset of those it can compute an attack slope from amplitude curves, which is the slope of the line from the beginning of the event to its peak. | The slope of an attack curve provides a relative measure of "alerting components," or immediately discriminable beginnings of a vocalization. |
| Roughness | `mir_roughness` | Mean, Med, StD, Range, Max, 1st Quart, 3rd Quart, IQR, Distance | A roughness value produced by computing the peaks of the audio spectrum and taking the average of the dissonance between all possible pairs of peaks; following Buyens et al. (2017), we reduce this to a single measure by taking the RMS-normalized mean. | Along with inharmonicity, roughness provides one measure of dissonance in a recording. Roughness similarly provides at least one measure of vocal clarity. |
| 85th Energy Percentile | `mir_rolloff85` | Whole | An estimate of the amount of high frequency in a signal measured by the frequency such that a 85% of the total energy is contained below it. | The 85th energy percentile allows a comparison of relative measures of high-frequency acoustics in a vocalization. |
| Inharmonicity | `mir_inharmonicity` | Whole | An estimate of the inharmonicity in the signal produced by identifying the number of partials that are not multiples of the fundamental frequency (i.e. those outside of the ideal harmonic range). | Along with roughness, inharmonicity provides a more precise measure of dissonance in a vocalization. |
| Tempo | `mir_tempo` | Whole | A tempo estimate made by detecting periodicities from MIR's event detection curves. Outputs a single number. | Tempo allows assessment of the speed or pace of a vocalization. |
| Pulse Clarity | `mir_pulseclarity` | Whole | Estimates the rhythmic clarity, or strength of the beats (Lartillot et al. 2008). | Pulse clarity provides a measure of the vocal clarity of a speaker or emphasis on individual utterances. |
| Rhythmic Variability | `npvi_total` | Recording | The nPVI equation measures the "average degree of durational contrast between adjacent events in a sequence" (Daniele & Patel, 2015). This makes it especially useful for comparing rhythmic units across language and music (i.e., syllables vs. notes). To automatically detect events, we used Mertens' (2004) syllable detection algorithm. | By providing a measure of durational contrast, nPVI_total is a measure of rhythmic complexity in a recording. |
| Rhythmic Variability | `npvi_phrase` | Phrase | In addition to detecting syllables, Mertens' algorithm detects phrases. Whereas npvi_total computes nPVI based on the whole file as a continuous phrase, this measure computes the nPVI for each detected phrase and reports the mean. In other words, it excludes the distances between the ends and beginnings of phrases. | nPVI_phrase provides a more granular measure of rhythmic complexity, within phrases, rather than between them. |
| Temporal Modulation | **tm_peak_hz** | Whole | The temporal modulation spectrum is the frequency decomposition of the amplitude envelope of a signal. This measures how loud something is at any given moment. We then measure how fast the loudness changes. For example: if someone sings a note every second, the spectrum will have a peak at 1Hz. If someone sings a note three times a second, but with an emphasis every three seconds, there will be a large peak at 1Hz, and a smaller peak at 3Hz. The peak of the spectrum is the frequency of the amplitude spectrum which has the highest root mean square of a given recording and represents a raw value of the recording's tempo. | The peak of the temporal modulation spectrum provides a measure of how maximally modulated, or variable, the onset of notes are in a recording, providing a raw measure of metre for speech and song. |
| Temporal Modulation | **tm_std_hz** | StD | The temporal modulation spectrum is the frequency decomposition of the amplitude envelope of a signal. This measures how loud something is at any given moment. We then measure how fast the loudness changes. For example: if someone sings a note every second, the spectrum will have a peak at 1Hz. If someone sings a note three times a second, but with an emphasis every three seconds, there will be a large peak at 1Hz, and a smaller peak at 3Hz. The standard deviation of the spectrum is taken as a measure of how exaggerated the peak is. | The standard deviation of temporal modulation allows for an assessment of the overall variability of temporal modulations in a recording, providing a coarse measure of rhythm, with a lower standard deviation leaning towards more monorhythmic signals. |
| Pitch | **praat_f0** | Mean, Med, StD, Range, Min, Max, 1st Quart, 3rd Quart, IQR | The fundamental frequency (f0) in Hertz for each recording | Pitch provides a fundamental measure of the highness or lowness, in frequency, of an utterance. Likewise, the shape of the pitch curve and the overall value of pitch is a common discriminable feature in both speech and song. |
| Pitch Space | **praat_f0travel** | Mean, Med, StD, Range, Max, 1st Quart, 3rd Quart, IQR | The distance between f0 at each .03125/sec interval to the next. | Pitch space provides a dynamic measure of pitch's range over time. |
| Pitch Rate | **praat_pitch_rate** | Whole, Med, IQR | The pitch rate is a measure of pitch change over time. In essence, the pitch rate provides a measure of pitch curve smoothness (a lower value corresponds to a smoother curve). | The pitch rate provides a measure of how smooth or variable pitch is over time. |
| Vowel Space | **praat_vowtrav** | Mean, Med, StD, Range, Max, 1st Quart, 3rd Quart, IQR | The Euclidian distance travelled in vowel space. This is equivalent to distance between the two formants. | Vowel space provides a measure of how much of the possible complex vowel space is used. |
| Vowel Space Travel Rate | **praat_vowtrav_rate** | Whole, Med, IQR | The Euclidian distance travelled in vowel space over a rate of time. This is equivalent to distance between two formants divided by rate of time. | Vowel travel rate provides a measure of how much of the vowel space is used over time, a relative measure of acoustic "flashiness" of a signal. |

| Label | Stub | Variables | Description | Significance |
| --- | --- | --- | --- | --- |
| Amplitude | `praat_intensity` | Mean, Med, StD, Range, Min, Max, 1st Quart, 3rd Quart, IQR, Distance | A measure of amplitude (loudness) in decibels | Amplitude provides a measure of how loud or quiet a vocalization is and can be compared between types within speakers |
| Amplitude Space | `praat_intensitytravel` | Mean, Med, StD, Range, Max, 1st Quart, 3rd Quart, IQR | The distance between amplitude at each .03125/sec interval to the next. | Intensity space provides a dynamic measure of intensity's range over time. |
| Amplitude Rate | `praat_intensity_rate` | Whole, Med, IQR | A measure of decay in intensity curves in each recording measured as change in amplitude over time. | The intensity rate provides a measure of how loud or soft amplitude changes over time. |
| 1st Formant | `praat_f1` | Mean, Med, StD, Range, Min, Max, 1st Quart, 3rd Quart, IQR | The frequency in Hertz of the 1st formant at each (.03125/sec) point | 1st formants are the 1st in a harmonic series following from the fundamental frequency and is important for a number of acoustic reasons. |
| Second Formant | `praat_f2` | Mean, Med, StD, Range, Min, Max, 1st Quart, 3rd Quart, IQR | The frequency in Hertz of the second formant at each (.03125/sec) point | Second formants are the second in a harmonic series following from the fundamental frequency, and along with the 1st formant, is used by listeners to perceive vowels. |
| File duration | `meta_length` |  | The length of the unedited sound files |  |
| Concatenated file duration | `meta_edit_length` |  | The length of the concatenated versions of the sound files |  |

**Extended Data Table 2.** The predictive influence of each of the acoustical features in distinguishing infant-directed from adult-directed vocalizations, chosen via two LASSO models (performance and the top six features for each model are depicted in Fig. 1b). The coefficients can be interpreted in a similar fashion to a logistic regression, i.e., as changes in the predicted log-odds ratio (with positive values indicating a higher likelihood of infant-directedness).

| Speech |  | Song |  |
| --- | --- | --- | --- |
| Acoustic feature | Coefficient | Acoustic feature | Coefficient |
| **Speech** |  | **Song** |  |
| Pitch (Median) | 2.449 | Vowel Travel (IQR) | 0.735 |
| Vowel Travel Rate (Median) | 0.677 | Intensity (Median) | -0.428 |
| Pitch (IQR) | 0.533 | Attack Curve Slope (Median) | -0.419 |
| Pulse Clarity | 0.231 | Roughness (Median) | -0.405 |
| Energy Roll-Off (85th %-ile) | -0.185 | Second Formant (IQR) | -0.285 |
| Second Formant (Median) | 0.170 | Energy Roll-Off (85th %-ile) | -0.255 |
| Roughness (IQR) | -0.167 | Inharmonicity | -0.171 |
| Attack Curve Slope (Median) | 0.152 | Attack Curve Slope (IQR) | 0.159 |
| Attack Curve Slope (IQR) | 0.119 | Pitch (IQR) | -0.156 |
| Inharmonicity | -0.073 | Vowel Travel Rate (IQR) | 0.117 |
| Tempo | -0.057 | Second Formant (Median) | -0.105 |
| Intensity (IQR) | 0.041 | Tempo | 0.080 |
|  |  | Pulse Clarity | 0.079 |
|  |  | Peak Tempo | 0.074 |
|  |  | Pitch (Median) | -0.042 |
|  |  | Rhythmic Variability (nPVI) | -0.028 |

**Extended Data Table 3.** Regression results from confirmatory analyses (corresponding with the boxplots in Fig. 2). The features tested here were limited to those with significant differences in the exploratory analyses. Statistics are from post-hoc linear combinations following multi-level mixed-effects models. Abbreviations: infant-directed (ID); adult-directed (AD).

| Comparison | Feature | Statistic | $\beta$ | $SE$ | $z$ | $p$ |
| --- | --- | --- | --- | --- | --- | --- |
| ID Speech vs. AD Speech |  |  |  |  |  |  |
|  | Intensity | Median | 0.081 | 0.052 | 1.542 | 0.123 |
|  | Acoustic Roughness | Median | -0.220 | 0.100 | -2.202 | 0.028 |
|  |  | IQR | -0.124 | 0.071 | -1.740 | 0.082 |
|  | Vowel Travel | IQR | 0.283 | 0.126 | 2.236 | 0.025 |
|  | Pitch (F<sub>0</sub>) | Median | 0.641 | 0.101 | 6.341 | <0.001 |
|  |  | IQR | 0.602 | 0.128 | 4.692 | <0.001 |
|  | Energy Roll-off (85 %ile) | Whole | -0.261 | 0.063 | -4.129 | <0.001 |
|  | Inharmonicity | Whole | -0.274 | 0.072 | -3.802 | <0.001 |
|  | Pulse Clarity | Whole | 0.213 | 0.069 | 3.092 | 0.002 |
|  | Vowel Travel Rate | Median | 0.514 | 0.116 | 4.412 | <0.001 |
|  |  | IQR | 0.519 | 0.123 | 4.234 | <0.001 |
| ID Song vs. AD Song |  |  |  |  |  |  |
|  | Intensity | Median | -0.138 | 0.048 | -2.905 | 0.004 |
|  | Acoustic Roughness | Median | -0.227 | 0.097 | -2.349 | 0.019 |
|  |  | IQR | -0.190 | 0.083 | -2.295 | 0.022 |
|  | Vowel Travel | IQR | 0.257 | 0.080 | 3.203 | 0.001 |
|  | Pitch (F<sub>0</sub>) | Median | -0.052 | 0.062 | -0.836 | 0.403 |
|  |  | IQR | -0.191 | 0.079 | -2.414 | 0.016 |
|  | Energy Roll-off (85 %ile) | Whole | -0.025 | 0.074 | -0.330 | 0.742 |
|  | Inharmonicity | Whole | -0.169 | 0.088 | -1.923 | 0.055 |
|  | Pulse Clarity | Whole | 0.064 | 0.111 | 0.579 | 0.562 |
|  | Vowel Travel Rate | Median | 0.179 | 0.094 | 1.896 | 0.058 |
|  |  | IQR | 0.211 | 0.088 | 2.396 | 0.017 |

**Extended Data Table 4.**
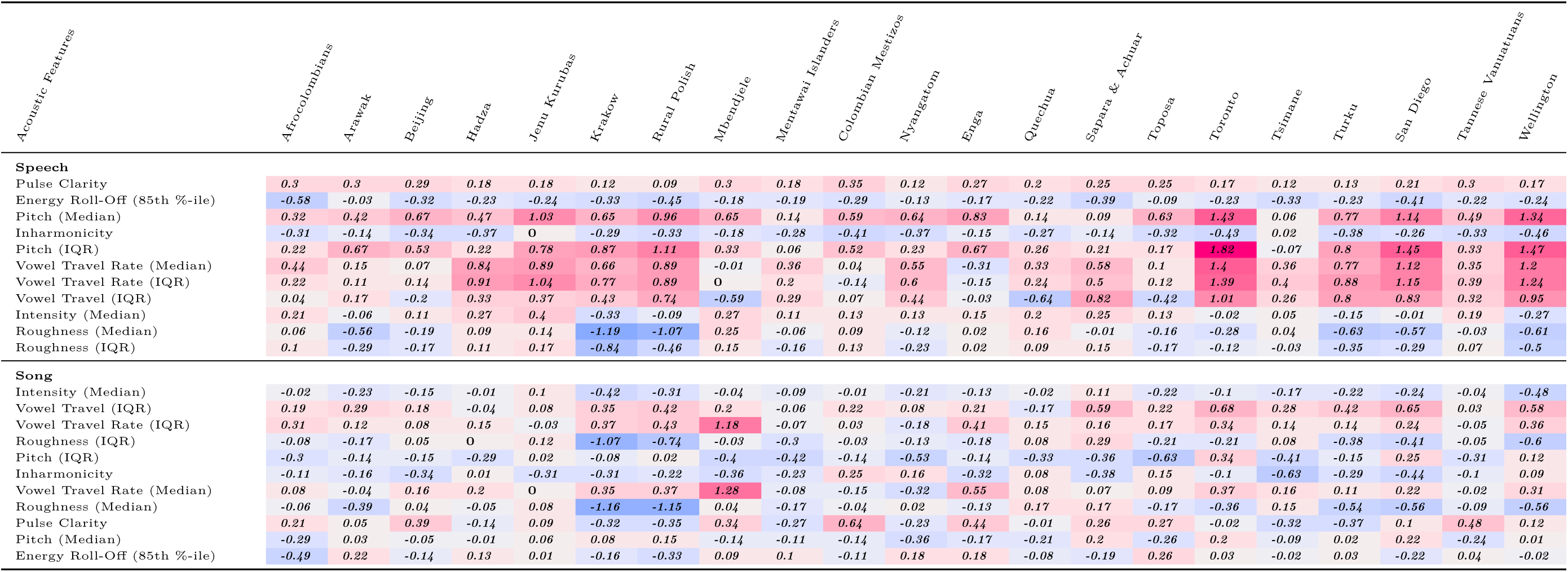
Estimated differences between infant-directed and adult-directed vocalizations, for acoustic feature, in each fieldsite (corresponding with the doughnut plots in Fig. 2). The estimates are derived from the random-effect components of the mixed-effects model reported in the Main Text. Cells of the table are shaded to facilitate the visibility of corpus-wide consistency (or inconsistency): redder cells represent features where infant-directed vocalizations have higher estimates than adult-directed vocalizations and bluer cells represent features with the reverse pattern. Within speech and song, acoustic features are ordered by their degree of cross-cultural regularity; some features showed the same direction of effect in all 21 societies (e.g., for speech, median pitch and pitch variability), whereas others were more variable.

**Extended Data Table 5.**
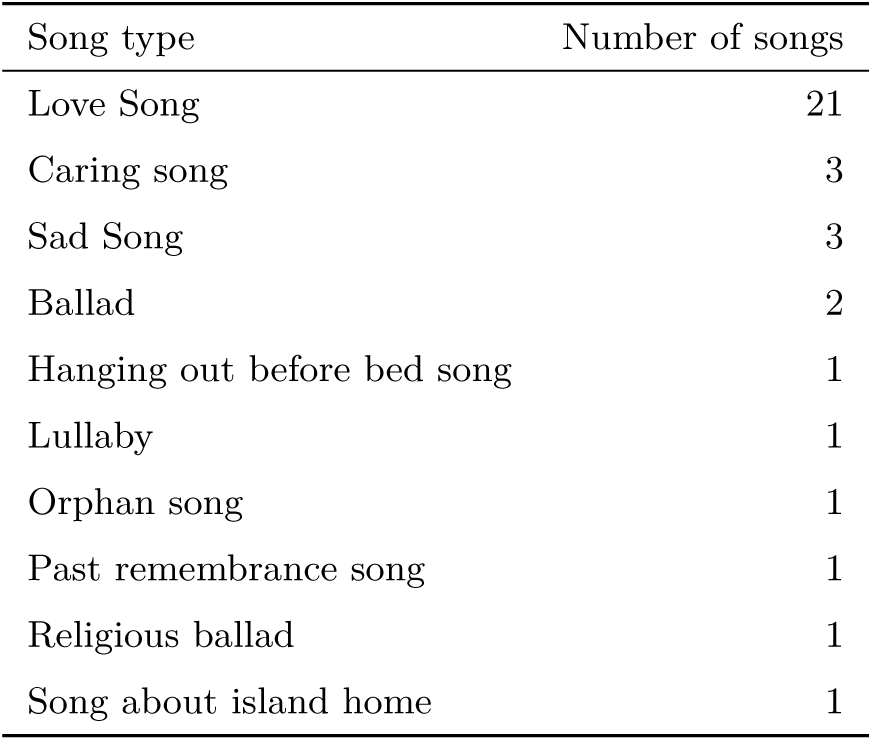
Adult-directed songs with descriptions rated as “soothing” by two independent annotators. A mixed-effects model estimating the difference in perceived infant-directedness across these vs. other adult-directed songs, adjusting for fieldsite- wise variability, found no statistically significant difference in responses (*b* = *−*0.011*, p* = .13).

| Song type | Number of songs |
| --- | --- |
| Love Song | 21 |
| Caring song | 3 |
| Sad Song | 3 |
| Ballad | 2 |
| Hanging out before bed song | 1 |
| Lullaby | 1 |
| Orphan song | 1 |
| Past remembrance song | 1 |
| Religious ballad | 1 |
| Song about island home | 1 |

**Extended Data Table 6.**
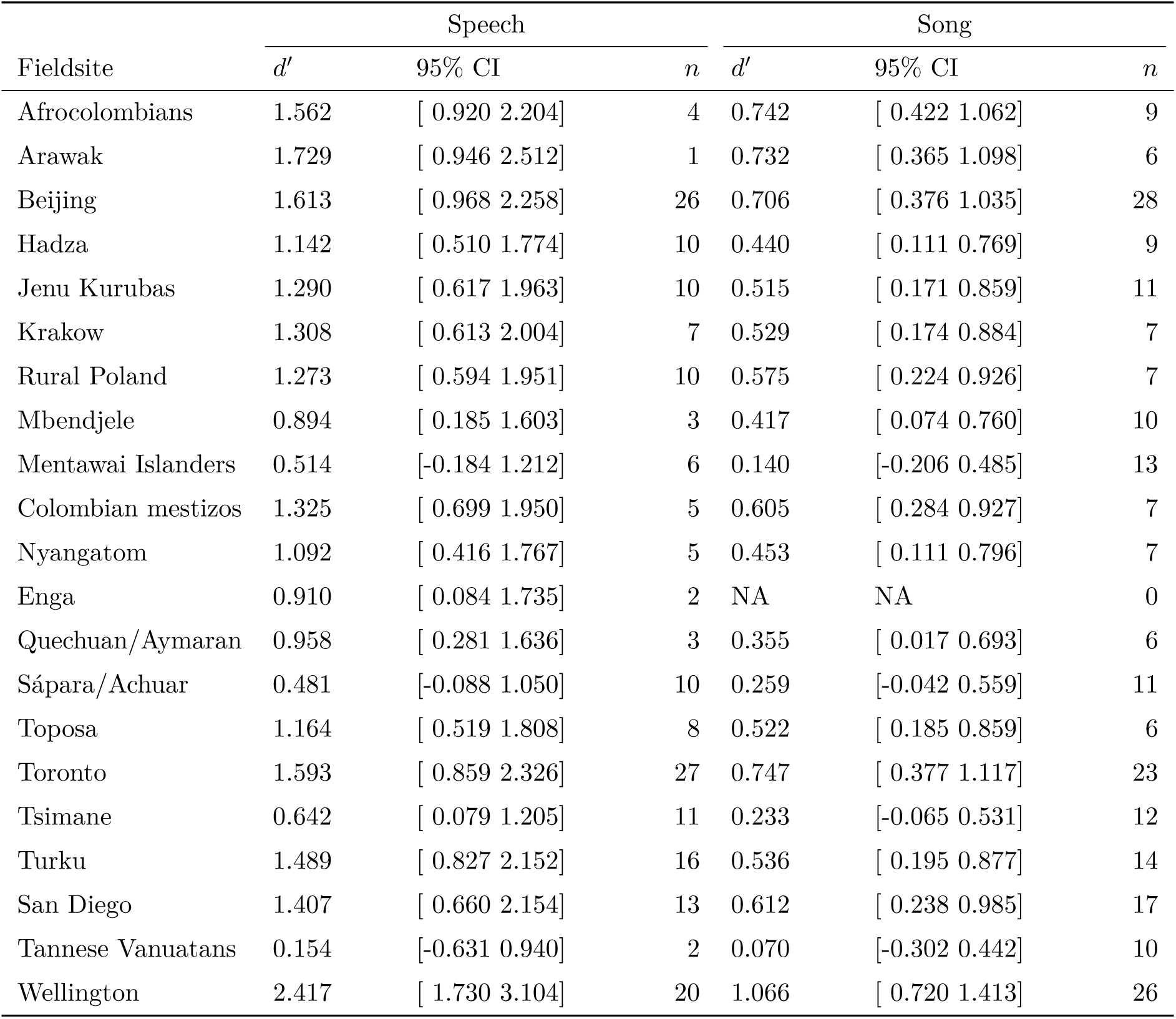
Estimated fieldsite-wise *d*-prime values, quantifying sensitivity to infant-directedness in speech and song, independent of response bias. Values are estimated as coefficients from mixed-effects model predicting *d^′^* from vocalization type, with random effects of fieldsite for each vocalization type. *n* refers to the number of vocalists that had a complete pair of vocalizations in the listener experiment (e.g., where one or both of the infant- and adult-directed vocalizations were not excluded due to confounds). Due to the strict exclusion procedure (see Methods), some fieldsites have very small samples, complicating the interpretation of these results, and one fieldsite had no observations for song. These exclusions only apply to the naïve listener experiment, however, and not the acoustic analyses reported elsewhere in this paper.

**Extended Data Table 7.** Demographics of United States participants. See notes and corresponding analyses in SI Text 1.5.

| Characteristic | % | N |
| --- | --- | --- |
| **Gender** |  |  |
| Female | 45.6% | 7352 |
| Male | 51.5% | 8299 |
| Other | 2.9% | 463 |
| [participant did not report] |  | 14 |
| **Ethnicity** |  |  |
| American Indian/Alaska Native | 1.4% | 207 |
| Asian | 23.3% | 3366 |
| Black or African-American | 3.7% | 536 |
| More than one race | 9.4% | 1351 |
| Native Hawaiian or other Pacific Islander | 0.9% | 131 |
| White | 61.2% | 8836 |
| [participant did not report] |  | 1701 |
| **Hispanic** |  |  |
| No | 87.6% | 12712 |
| Yes | 12.4% | 1804 |
| [participant did not report] |  | 1612 |
| **Annual household income** |  |  |
| Under \$10,000 | 9.1% | 912 |
| \$10,000 to \$19,999 | 8.8% | 879 |
| \$20,000 to \$29,999 | 7.4% | 747 |
| \$30,000 to \$39,999 | 7.5% | 755 |
| \$40,000 to \$49,999 | 7.4% | 747 |
| \$50,000 to \$74,999 | 14.7% | 1471 |
| \$75,000 to \$99,999 | 12.2% | 1227 |
| \$100,000 to \$150,000 | 17.9% | 1795 |
| Over \$150,000 | 15.0% | 1503 |
| [participant did not report] |  | 6092 |

**Extended Data Table 8.** Factor loadings for the top three principal components reported in Extended Data Fig. 3.

| Principal Component 1 |  | Principal Component 2 |  | Principal Component 3 |  |
| --- | --- | --- | --- | --- | --- |
| Feature | Weighting | Feature | Weighting | Feature | Weighting |
| Amplitude Space (Mean) | -0.201 | Amplitude (Mean) | 0.269 | Pitch (Mean) | -0.309 |
| Amplitude Space Travel Rate (Median) | -0.200 | Amplitude (Median) | 0.264 | Pitch (3rd Quartile) | -0.304 |
| Pitch Space_rate (IQR) | -0.199 | Amplitude (3rd Quartile) | 0.261 | Pitch (Median) | -0.295 |
| Pitch Space Travel Rate (Whole) | -0.198 | Amplitude (1st Quartile) | 0.241 | Pitch (1st Quartile) | -0.252 |
| Amplitude Space Travel Rate (IQR) | -0.195 | Roughness (3rd Quartile) | 0.215 | Pitch (IQR) | -0.222 |
| Amplitude Space (Median) | -0.188 | Roughness (IQR) | 0.214 | Roughness (1st Quartile) | 0.213 |
| Amplitude Space Travel Rate (Whole) | -0.188 | Roughness (Standard Deviation) | 0.200 | Roughness (Median) | 0.192 |
| Pitch Space (3rd Quartile) | -0.187 | Roughness (Median) | 0.191 | Pitch (Standard Deviation) | -0.174 |
| Pitch Space (IQR) | -0.186 | Amplitude (Maximum) | 0.183 | Roughness (3rd Quartile) | 0.151 |
| Amplitude Space (1st Quartile) | -0.186 | Roughness (Range) | 0.169 | Roughness (IQR) | 0.146 |
| Amplitude Space (3rd Quartile) | -0.185 | Roughness (Maximum) | 0.169 | Roughness (Mean) | 0.143 |
| Vowel Space Travel Rate (Median) | -0.185 | Amplitude (Minumum) | 0.168 | Amplitude Space (Range) | -0.143 |
| Pitch Space (Mean) | -0.181 | Roughness (Mean) | 0.155 | Amplitude Space (Maximum) | -0.143 |
| Vowel Space Travel Rate (IQR) | -0.180 | 1st Formant (1st Quartile) | 0.150 | Pitch Space (1st Quartile) | -0.130 |
| Amplitude Space (IQR) | -0.179 | Roughness (1st Quartile) | 0.136 | Amplitude (3rd Quartile) | -0.127 |
| Vowel Space Travel Rate (Whole) | -0.178 | 1st Formant (Standard Deviation) | -0.133 | Pitch (Maximum) | -0.123 |
| Pitch Space_rate (Median) | -0.177 | Amplitude Space (Maximum) | 0.132 | Amplitude (Mean) | -0.122 |
| Vowel Space (Mean) | -0.170 | Amplitude Space (Range) | 0.132 | Amplitude (Median) | -0.121 |
| Amplitude Space (Standard Deviation) | -0.161 | Vowel Space (IQR) | -0.129 | 85th Energy Percentile | 0.120 |
| Vowel Space (Median) | -0.161 | Second Formant (Mean) | -0.126 | Second Formant (Minumum) | 0.114 |
| Vowel Space (Standard Deviation) | -0.159 | Vowel Space (3rd Quartile) | -0.126 | Amplitude (Maximum) | -0.109 |
| Vowel Space (1st Quartile) | -0.159 | 1st Formant (Minumum) | 0.125 | Amplitude (1st Quartile) | -0.109 |
| Vowel Space (3rd Quartile) | -0.151 | Second Formant (3rd Quartile) | -0.124 | Second Formant (IQR) | -0.109 |
| Pitch Space (Standard Deviation) | -0.151 | 1st Formant (Range) | -0.123 | Inharmonicity | 0.108 |
| Pitch Space (Median) | -0.151 | Second Formant (Median) | -0.120 | Pitch Space (Maximum) | -0.106 |
| Vowel Space (IQR) | -0.143 | Second Formant (Maximum) | -0.119 | Pitch Space (Range) | -0.106 |
| Amplitude (IQR) | -0.126 | Second Formant (Range) | -0.117 | Amplitude (Range) | -0.105 |
| Temporal Modulation (Peak) | -0.107 | 1st Formant (Median) | 0.114 | 1st Formant (Mean) | 0.103 |
| nPVI Recording | 0.100 | Vowel Space (Mean) | -0.111 | Temporal Modulation (Standard Deviation) | -0.103 |
| Amplitude (Standard Deviation) | -0.098 | 1st Formant (Maximum) | -0.108 | Second Formant (Standard Deviation) | -0.102 |

